# Harmonin homology domain-mediated interaction of RTEL1 helicase with RPA and DNA provides mechanistic insight into its role in DNA repair

**DOI:** 10.1101/2022.08.08.503141

**Authors:** Niranjan Kumar, Arushi Taneja, Meenakshi Ghosh, Ulli Rothweiler, Nagalingam Ravi Sundaresan, Mahavir Singh

**Affiliations:** Molecular Biophysics Unit, Indian Institute of Science, Bengaluru, 560012, India; Department of Microbiology and Cell Biology, Indian Institute of Science, Bengaluru, 560012, India; The Norwegian Structural Biology Centre, Department of Chemistry, The Arctic University of Norway, N-9037 Tromsø, Norway

**Keywords:** RTEL1, RPA, Harmonin homology domain, DNA repair, NMR chemical shift perturbations, X-ray crystal structure, D-loop

## Abstract

The regulator of telomere elongation helicase 1 (RTEL1) is an Fe-S cluster containing helicase that plays important roles in telomere DNA maintenance, DNA repair, and genome stability. It is a modular protein comprising a helicase domain, two tandem harmonin homology domains 1 & 2 (HHD1 and HHD2), and a Zn^2+^ binding RING domain. In this study, we have unravelled a novel interaction between RTEL1 and replication protein A (RPA) and shown their co-localization upon DNA damage in the cells. Using NMR spectroscopy, we show that 32C domain of RPA and DNA competitively bind with HHD2 of RTEL1. To understand the structural basis of HHD2 – 32C and HHD2 - DNA interactions, we have determined a 1.6 Å resolution crystal structure of HHD2. NMR chemical shift perturbations-based mapping revealed the 32C and DNA binding surface on HHD2 of RTEL1. Together, these results establish an interplay among RTEL1, RPA, and DNA that provide mechanistic insights into the RTEL1 recruitment at DNA during the processes of replication, repair, and recombination.

## INTRODUCTION

The regulator of telomere elongation helicase 1 (RTEL1) is an Fe-S cluster and DEAH motif-containing DNA helicase with ATP-dependent 5’ to 3’ helicase activity (Estep and Brosh, 2018; Uringa et al., 2011). RTEL1 plays essential roles in telomere maintenance, DNA repair, meiotic recombination, and genome-wide replication (Sarek et al., 2015; Uringa et al., 2012; Uringa et al., 2011; Vannier et al., 2012; Vannier et al., 2014). It is a modular protein consisting of an N-terminal helicase domain followed by two tandem harmonin homology domains 1 & 2 (HHD1 and HHD2), a PCNA-interacting protein-box (PIP-box) motif, and a C-terminal C4C4 type RING domain (Vannier et al., 2014) (Figure 1A). RTEL1 is an anti-recombinase and executes non-crossovers by promoting the synthesis-dependent strand annealing (SDSA) pathway of homologous recombination through disassembling D-loop intermediates during the DNA repair and meiotic recombination processes (Barber et al., 2008; Uringa et al., 2011). RTEL1 interacts with the replisome’s proliferating cell nuclear antigen (PCNA) through its PIP-box motif and helps genome-wide replication (Sarek et al., 2015; Vannier et al., 2013). The central HHD1 and HHD2 are predicted to mediate protein-protein interactions (Faure et al., 2014). Several mutations in RTEL1 are associated with genetic diseases Hoyeraal-Hreidarsson syndrome (HHS) and familial pulmonary fibrosis (FPF) (Deng et al., 2013; Jullien et al., 2016; Stuart et al., 2015; Vannier et al., 2014; Walne et al., 2013). These diseases are associated with short telomere length in patients resulting in premature aging, bone marrow failure, and predisposition to cancer.

**Figure 1.**
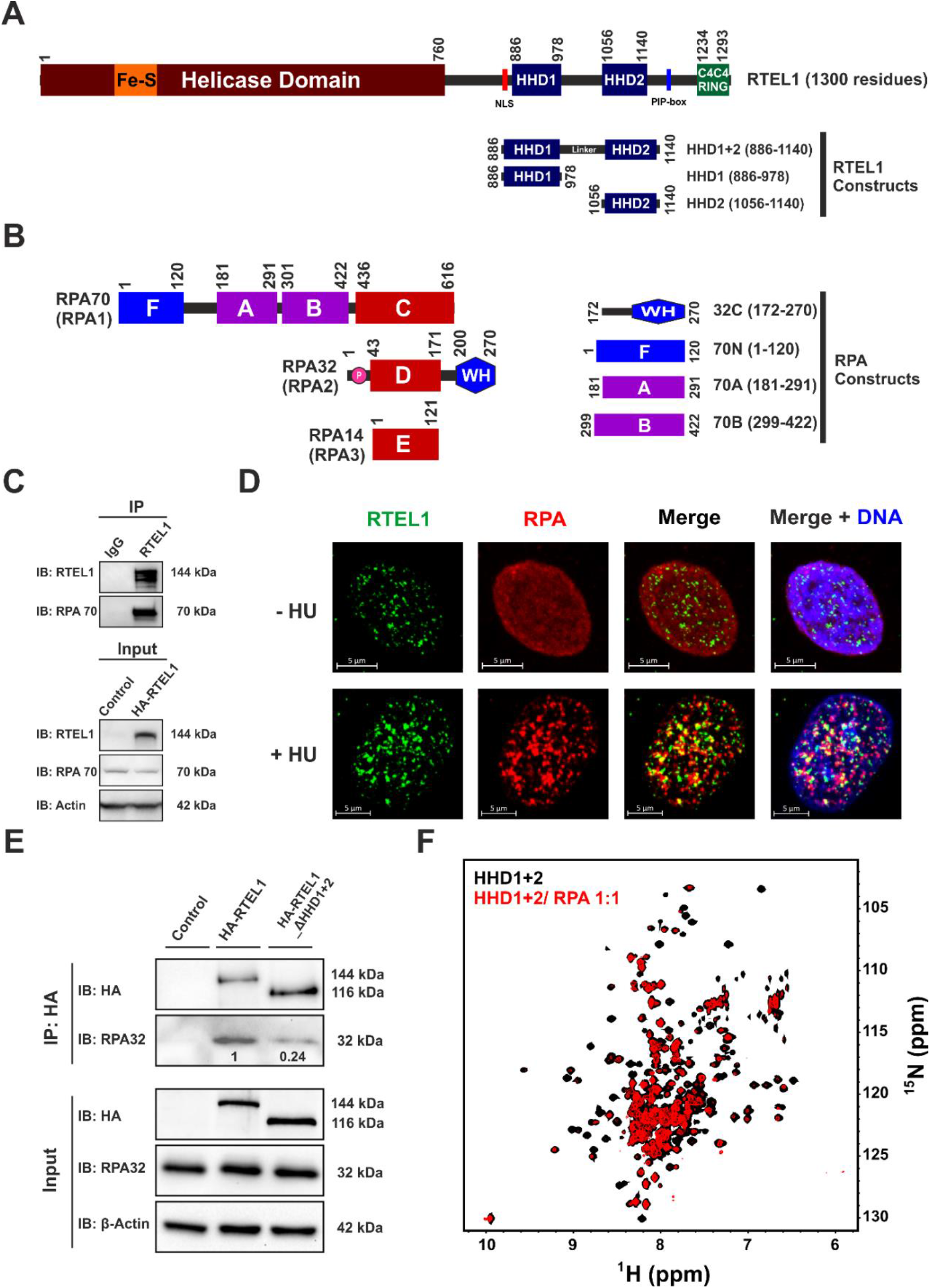
RTEL1 directly interacts with RPA. **(A)** Domain organization of human RTEL1. HHD constructs used in this work are shown with domain boundaries. **(B)** Domain organization of human RPA trimeric complex (consisting of RPA70, RPA32, and RPA14). Various constructs of RPA used in this study are shown with their domain boundaries. **(C)** Whole-cell extracts of HEK293T cells overexpressing HA-tagged human RTEL1 were immunoprecipitated (IP) with mentioned antibodies. The immunocomplexes (*upper panel*) and input (*bottom panel*) were analyzed through immunoblotting (IB) with indicated antibodies. **(D)** Representative immunofluorescence images showing co-localization of endogenous RTEL1 and RPA in the nucleus of HeLa cells. Cells were untreated (*upper panel*) or treated (*bottom panel*) with 4 mM Hydroxyurea (HU) for 48 h, paraformaldehyde-fixed, and immuno-stained for RTEL1 (green) and RPA (red). Nuclear DNA (blue) was counterstained with Hoechst 33342. **(E)** Representative immunoblot showing co-immunoprecipitation of RPA with HA-RTEL1 and HA-RTEL1_ΔHHD1+2. Whole-cell extracts of HEK293T cells, overexpressing HA-tagged human RTEL1 and its deletion construct RTEL1_ΔHHD1+2, were immunoprecipitated (IP) using an anti-HA antibody. The immunocomplexes (*upper panel*) and input (*bottom panel*) were analyzed through immunoblotting (IB) with indicated antibodies. RTEL1 and RPA interaction reduced by about 75% upon deletion of the HHD1+2 domain. Data were analyzed from two independent biological experiments using ImageJ software. The normalized value of RPA is mentioned below the respective bands. **(F)** Overlay of ^1^H-^15^N TROSY HSQC spectra of ^15^N-labelled HHD1+2 in the absence (black) and presence (red) of RPA complex (at 1:1 molar ratio).

HHDs are the most recent entrant to newly classified αα-hub domain-containing proteins (Colcombet-Cazenave et al., 2021). So far, nine HHDs have been identified in six proteins, viz. Harmonin, Whirlin, Delphilin, PDZD7, CCM2, and RTEL1 (Bugge et al., 2021; Colcombet-Cazenave et al., 2021; Staby et al., 2021). The HHD1 and HHD2 of RTEL1 harbor several mutations associated with HHS and FPF, underscoring their functional importance (Faure et al., 2014). In a recent study, a region encompassing the HHD1 of RTEL1 was shown to interact with SLX4 (Takedachi et al., 2020; Vannier et al., 2012; Wu et al., 2020).

Replication protein A (RPA) is a trimeric protein complex (consisting of RPA70, RPA32, and RPA14 subunits) (Figure 1B) that binds single-stranded DNA (ssDNA) with high affinity and sequence-independent manner (Wold, 1997). ssDNA is an intermediate product in the replication, recombination, and repair (3R) pathways (Branzei and Szakal, 2021). Since RPA and many DNA helicases (including RTEL1) are critical players in the 3R pathways, they play interactive roles in maintaining genomic integrity (Awate and Brosh, 2017). Many DNA helicases like BLM, WRN, and FANCJ have been shown to interact physically and functionally with RPA, but the direct RTEL1 – RPA interaction has remained uncharacterized so far (Awate and Brosh, 2017; Caldwell and Spies, 2020; Jullien et al., 2016). Homologous recombination is an essential cellular process for the accurate repair of DNA double-strand breaks (DSBs), meiotic recombination, and the restart of stalled replication forks. The displacement loop (D-loop) is formed as an intermediate structure during homologous recombination. The D-loop is also present within the t-loop structure of telomere DNA (de Lange, 2004). TRF2-mediated recruitment of RTEL1 at the telomere promote t-loop unwinding (Sarek et al., 2015). In an *in vitro* assay, RTEL1 was shown to preferentially dissociate the D-loops with 3’ invasion (Youds et al., 2010). However, the mechanism through which RTEL1 is recruited to the non-telomeric D-loop for its proper disassembly during DNA repair and meiotic recombination events was undefined.

Here, we report that human RTEL1 directly interacts and co-localizes with RPA upon DNA damage. Interestingly, this interaction is mediated by the tandem harmonin homology domains of RTEL1. Using NMR chemical shift perturbation experiments, we showed that the HHD2 of RTEL1 interacts exclusively with the winged-helix domain 32C of RPA. We have characterized the HHDs using NMR spectroscopy and solved the X-ray crystal structure of HHD2 of human RTEL1. The structure of HHD2 revealed a unique surface charge potential with positive and negative surfaces. Using NMR spectroscopy and isothermal titration calorimetry (ITC), we have shown that the HHD2 also interacts with DNA using the RPA 32C binding surface. This study established HHD2 as a novel accessory domain of RTEL1 that mediates both protein-protein and protein-DNA interactions. Interestingly, we have also shown that ssDNA competitively displaces the RPA 32C from RTEL1 HHD2 – RPA 32C complex. The interplay among RTEL1, RPA, and DNA suggests a possible mechanism of RPA-mediated recruitment of RTEL1 at D-loop present at the DNA repair and recombination sites.

## RESULTS

### RTEL1 physically and functionally interacts with RPA

As per the BioGRID database (Oughtred et al., 2021), RTEL1 is included in the interactome of the RPA-ssDNA complex (Maréchal et al., 2014). However, there is no report of direct interaction between RTEL1 and RPA. Both RTEL1 and RPA are involved in several DNA metabolic pathways. Therefore, we investigated for any possible physical and functional interaction between RTEL1 and RPA. First, the HEK293T cells were transfected with pcDNA3.1-N-HA-RTEL1 for transient overexpression of HA-tagged RTEL1. A co-immunoprecipitation experiment was performed, and the results showed that endogenous RPA co-immunoprecipitated with RTEL1, suggesting physical interaction between RPA and RTEL1 (Figure 1C). Since RPA and RTEL1 have essential roles in the DNA repair pathways, we treated HeLa cells with hydroxyurea (a DNA damaging agent) and performed an immunofluorescence microscopy-based experiment. Upon DNA damage, a subset of endogenous RTEL1 and RPA foci co-localize (Figure 1D), thus indicating the functional nature of RTEL1-RPA interaction.

Analysis of the RPA interaction sites on the BLM, WRN, and FANCJ helicases indicates that a motif/domain other than the helicase domain interacts with the RPA (Figure S1A) (Awate and Brosh, 2017; Kang et al., 2018; Yeom et al., 2019). Since the HHDs in RTEL1 were classified as putative protein-protein interaction domains (Colcombet-Cazenave et al., 2021; Faure et al., 2014), we hypothesised that HHDs could harbor potential RPA binding sites (Figure S1A). To confirm this hypothesis, we overexpressed the HHD1+2 (HHD1-linker-HHD2) deletion construct of RTEL1 (pcDNA3.1-N-HA-RTEL1-ΔHHD1+2) in HEK293T cells. Deletion of HHD1+2 resulted in a significant reduction of co-immunoprecipitated RPA (Figure 1E), suggesting that, indeed, HHDs of RTEL1 interact with RPA.

Protein-protein interactions in the cell are often weak and transient. NMR chemical shifts are extremely sensitive to changes in the environment. Therefore, weak but biologically relevant interactions can be studied using NMR chemical shift perturbations (Vaynberg and Qin, 2006). The ^15^N-labeled HHD1+2 was titrated with human RPA. We observed resonance peak broadening and chemical shift perturbations (CSP) of several residues, which clearly indicate a direct interaction between HHD1+2 and RPA (Figure 1F, Figure S1B and S1D).

### The tandem harmonin homology domains of RTEL1 are connected through an intrinsically disordered linker region

Although predicted to be consisting of a harmonin-N-like fold using the hydrophobic cluster analysis (Faure et al., 2014), HHD1 and HHD2 of the RTEL1 have not been characterized experimentally for their structures. Bioinformatic analysis of the tandem harmonin homology domains of RTEL1 indicates that a disordered linker region separates HHD1 and HHD2 (Figure 2A). We have purified the ^15^N-labeled HHD1, HHD2, and HHD1+2 (Figure S1B) and recorded the ^1^H-^15^N 2D HSQC spectrum for each protein. The spectra of HHD1 and HHD2 overlaid well on the spectrum of HHD1+2 (Figure 2B). The extra cross-peaks arising from the linker region clustered at the centre of the spectrum (∼8.2 ppm in the ^1^H dimension) of HHD1+2, indicating the disordered nature of the linker. Moreover, no significant chemical shift perturbations were observed in the 2D ^1^H-^15^N HSQC spectrum of HHD2 when titrated with the unlabelled HHD1 (Figure S1C). Altogether, these results suggest that individual HHD1 and HHD2 are independently folded domains separated by a disordered linker region. We have recently reported near-complete backbone and sidechain resonance assignments of HHD1 and HHD2 (Kumar et al., 2022).

**Figure 2.**
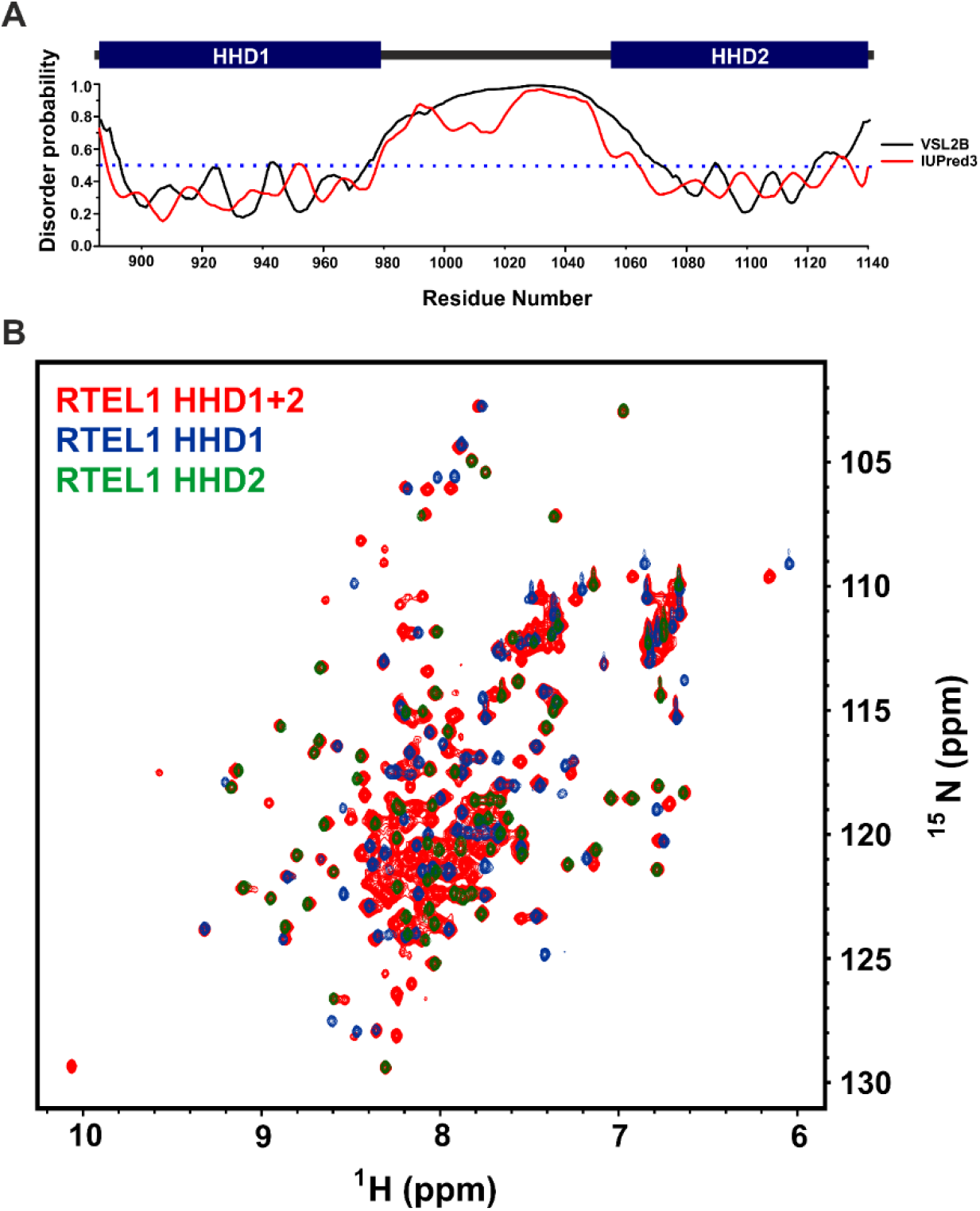
Intrinsically disordered linker connects HHD1 and HHD2 of RTEL1. **(A)** Prediction of the intrinsically disordered region in the RTEL1 HHD1+2 domain using bioinformatic tools VSL2B and IUPred3. The blue dash line indicates the cut-off value (0.5) for disorder probability. **(B)** Overlay of ^1^H-^15^N HSQC spectra of HHD1 (blue), HHD2 (green), and HHD1+2 (red).

### RPA interacts with HHD2 of RTEL1 through its winged-helix domain 32C

RPA is a heterotrimeric multi-domain protein complex consisting of three subunits, RPA70, RPA32, and RPA14. The largest subunit of RPA, i.e., RPA70 (RPA1), consists of four OB-fold domains (70N, 70A, 70B, and 70C domains), RPA32 (RPA2) subunit consists of OB-fold 32D and winged-helix 32C domains, whereas the RPA14 (RPA3) subunit consists of OB-fold domain 14E (Figure 1B). Protein and DNA binding roles have been assigned to different domains of RPA. The 70N, 70A, 70B, and 32C domains of RPA interact with partner proteins (Caldwell and Spies, 2020). We took an NMR spectroscopy-based titration approach to map the interaction between HHDs of RTEL1 and different domains of RPA. Uniformly ^15^N-labeled samples of 70N, 70A, 70B, and 32C domains of RPA (Figure S1D) were titrated with the HHD1+2 domain of RTEL1. No significant CSPs were observed in the spectra of 70N, 70A, and 70B (Figure S2A-2C), while we observed distinct CSPs in the spectrum of 32C upon titration with HHD1+2 (Figure S2D). The reverse titrations (^15^N-labelled HHD1+2 titrated with RPA 32C) also showed distinct CSPs in the spectrum of HHD1+2 (Figure S2E).

In subsequent experiments, individual HHD1 and HHD2 were titrated with RPA 32C. No significant CSPs were observed in the spectrum of HHD1 (Figure S2F); however, distinct CSPs of several residues were observed in the case of HHD2 (Figures 3A and 3B). We also performed a reverse NMR titration of ^15^N-labelled 32C with HHD2 (Figure 3C and 3D). Again, we observed distinct CSPs in several residues of 32C. Interestingly, these are the same set of residues of 32C, which were perturbed upon titration with HHD1+2 (Figure S2D). Based on these results, we conclude that the HHD2 domain of RTEL1 primarily interacts with the 32C domain of RPA.

**Figure 3.**
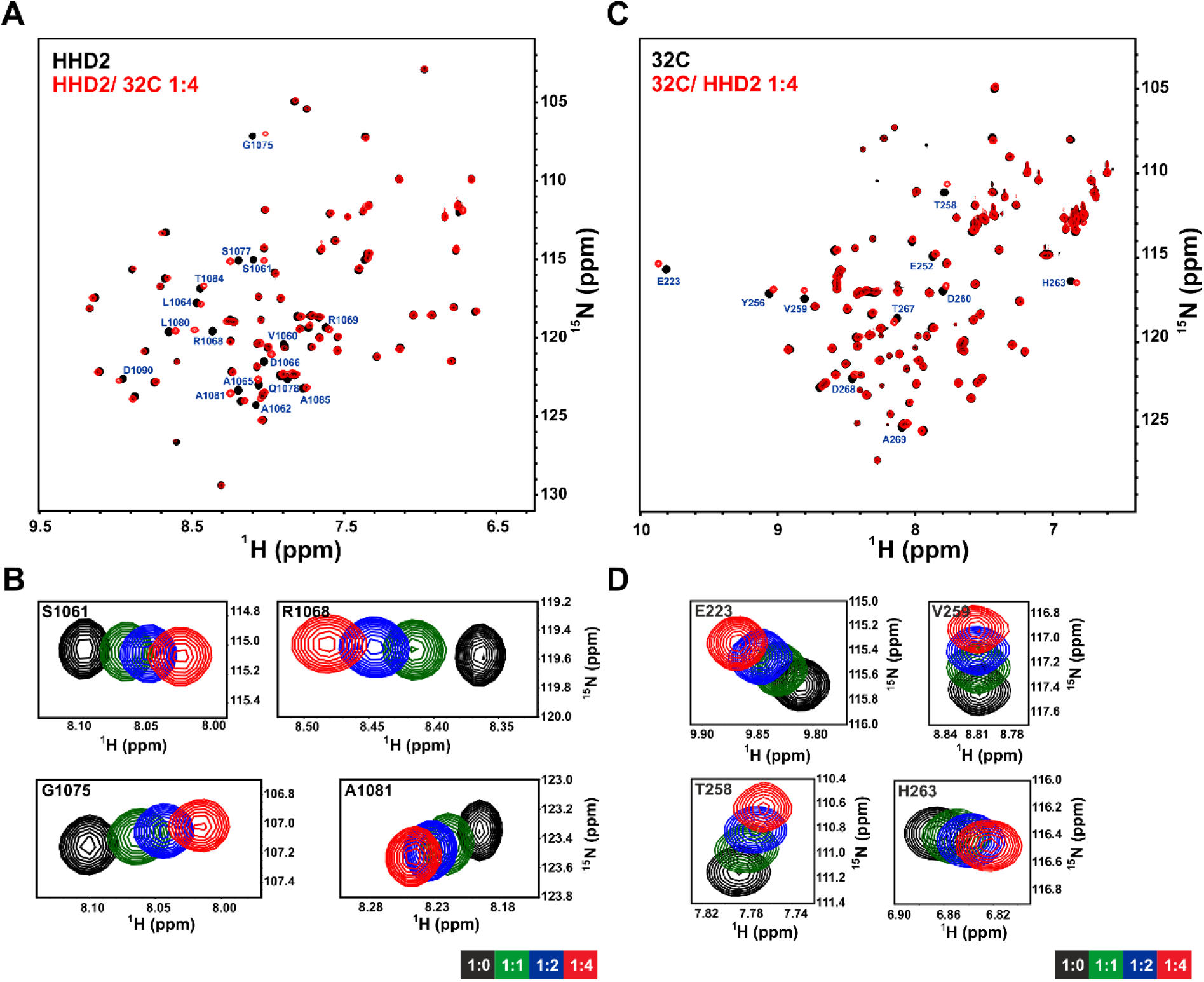
NMR titration of RTEL1 HHD2 and RPA 32C domains. **(A)** Overlay of ^1^H-^15^N HSQC spectra of ^15^N-labelled HHD2 in the absence (black) and presence (red) of RPA 32C (at 1:4 molar ratio). Residues that showed large CSPs are labeled. **(B)** ^1^H-^15^N cross-peaks trajectory of representative residues S1061, R1068, G1075, and A1081 of HHD2 upon titration with RPA 32C at indicated molar ratios. **(C)** Overlay of ^1^H -^15^N HSQC spectra of ^15^N-labelled 32C in the absence (black) and presence (red) of HHD2 (at 1:4 molar ratio). Residues that showed large CSPs are labeled. **(D)** ^1^H -^15^N cross-peaks trajectory of representative residues E223, T258, V259, and H263 of 32C upon titration with HHD2 at indicated molar ratios.

### Structural basis of RTEL1 HHD2 and RPA 32C interaction

To understand the structural basis of RPA 32C and RTEL1 HHD2 interaction, we solved the structure of HHD2 at a resolution of 1.6 Å (Table 1) using X-ray crystallography (details in Methods section). The structure revealed that HHD2 consists of five helices, H1-H5, connected by a short loop. All the helices are mainly α-helical (3.6_13_) except the N-and C-termini of H4 and the N-terminal of H5, which consists of short 3_10_ helices (Figure 4A and 4B). The secondary structures match well with the NMR chemical shift indexing-based secondary structures (Kumar et al., 2022). The overall topology of RTEL1 HHD2 is similar to the N-terminal domain of the human Harmonin (Faure et al., 2014). Evolutionarily conserved residues are mainly present in the core region and stabilize the tertiary structure (Figure 4A-4C). The electrostatic surface potential of HHD2 revealed a distinct pattern of positively and negatively charged surfaces (Figure 4D).

**Table 1.** Crystallographic data collection and structure refinement statistics of RTEL1 HHD2.

|  |  |
| --- | --- |
| PDB ID | 7WU8 |
| **Integration** |  |
| Space group | P 1 2<sub>1</sub> 1 |
| Unit Cell constants | a = 65.64 Å, b = 60.96 Å, c = 79.71 Å<br>α = 90.0°, β = 95.64°, γ = 90.0° |
| Wavelength (Å) | 0.9763 |
| Resolution range (Å) | 48.16 – 1.60 (1.70-1.60) |
| Observed reflections | 278293 (42947) |
| Unique reflections | 80150 (12698) |
| Data completeness (%) | 96.8 (95.3) |
| < I/σ(I) > | 1.56 (at 1.6 Å) |
| R<sub>meas</sub> (%) | 5.3 (64.0) |
| CC (1/2) | 99.9 (80.2) |
| **Refinement** |  |
| Number of reflections | 78046 |
| R<sub>work</sub>, R<sub>free</sub> | 0.182, 0.211 |
| No. of molecules in A.S.U. | 7 |
| Total number of atoms | Total: 5128,<br>Solvent: 734, Non-solvent: 4394 |
| Average B, all atoms (Å<sup>2</sup>) | 30.0 |
| R.M.S.D<br>bond length(Å) / Angles (°) | 0.015/ 1.895 |
| Ramachandran outliers<br>Favored/allowed/outliers (%) | 100/0/0 |
| Clash score | 3 |
| Molprobity score | 1.01 (100<sup>th</sup> percentile) |
| *Highest resolution shell is shown in parentheses |  |

**Figure 4.**
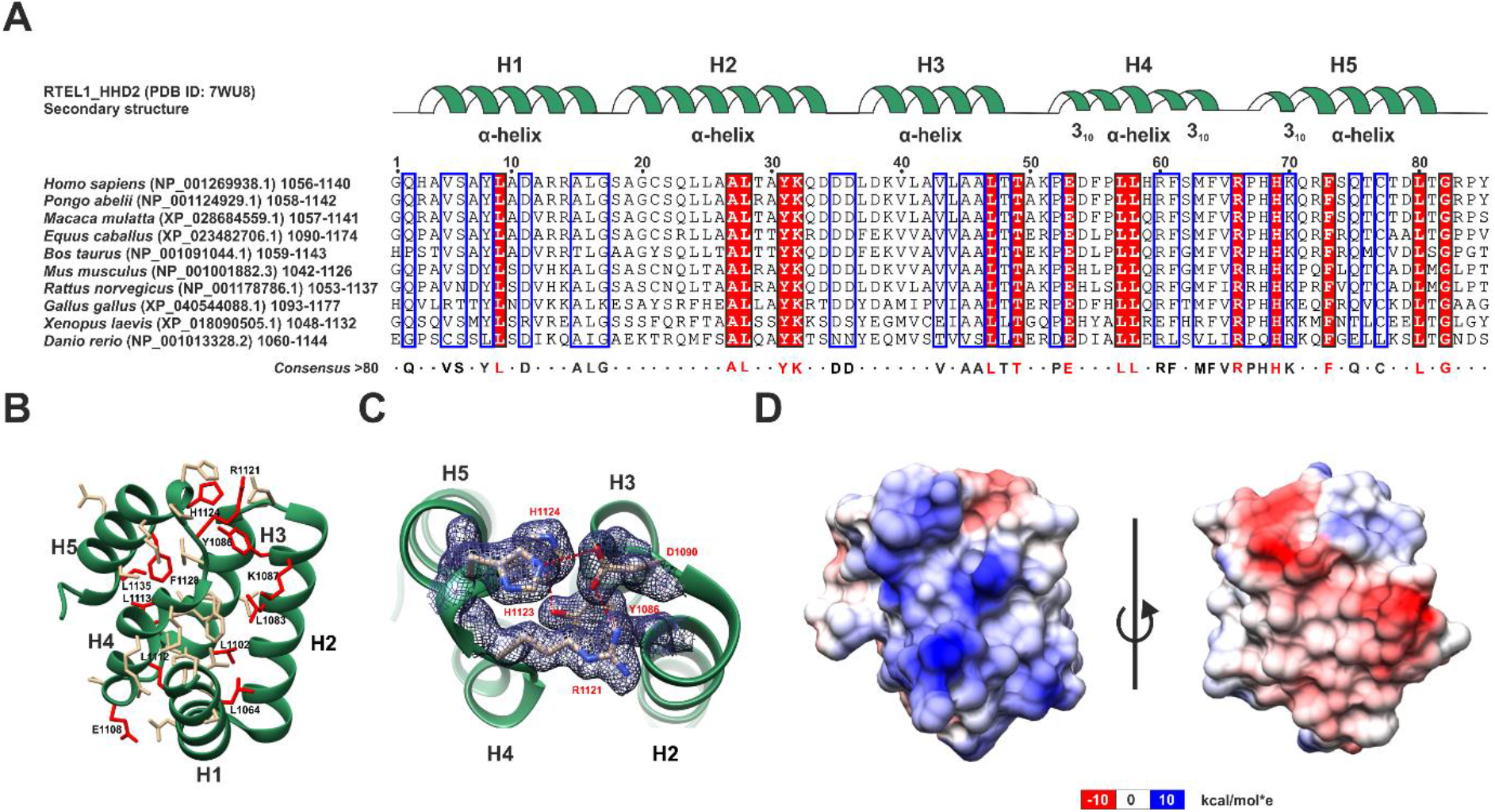
Structure of HHD2 domain of RTEL1. **(A)** Multiple sequence alignment of HHD2 of RTEL1 proteins from different vertebrate species. The secondary structure corresponding to the HHD2 of human RTEL1 is depicted at the top (green ribbon). The consensus sequence, including the conserved residues (red), is shown at the bottom. **(B)** 1.6 Å crystal structure of HHD2 domain of human RTEL1. Sidechains of evolutionarily conserved residues are shown as stick models (fully conserved residues and partially conserved residues are shown in red and gold, respectively). **(C)** Electron density map of a representative region consisting of conserved residues Y1086, D1090, R1121, H1123, and H1124. One salt bridge (D1090 OD1 – R1121 NH1) and two H-bonds (Y1086 OH – H1124 ND1 and D1090 OD2 – H1123 NE2) are shown (red dash line). **(D)** Unique surface electrostatic potential map of HHD2. One surface (consisting of H1 and H2 helices) of the protein is positively charged (blue), while the other surface (consisting of H2, H3, and H5) is negatively charged (red).

Our recently deposited chemical shifts of HHD2 (BMRB entry 51077) (Kumar et al., 2022) were used to map the RPA 32C binding surface on the HHD2 crystal structure. The observed CSPs in the titration of ^15^N-labelled HHD2 with 32C (Figure 3A) were calculated and plotted (Figure 5A). The HHD2 residues (A1059, V1060, S1061, A1062, Y1063, L1064, A1065, D1066, A1067, R1068, R1069, G1075, S1077, Q1078, L1079, L1080, A1081, A1082, T1084, K1087, D1090, and D1134) that showed perturbation more than the average CSPs upon 32C binding were mainly present in the N-terminal helices H1 and H2 (Figure 5B).

**Figure 5.**
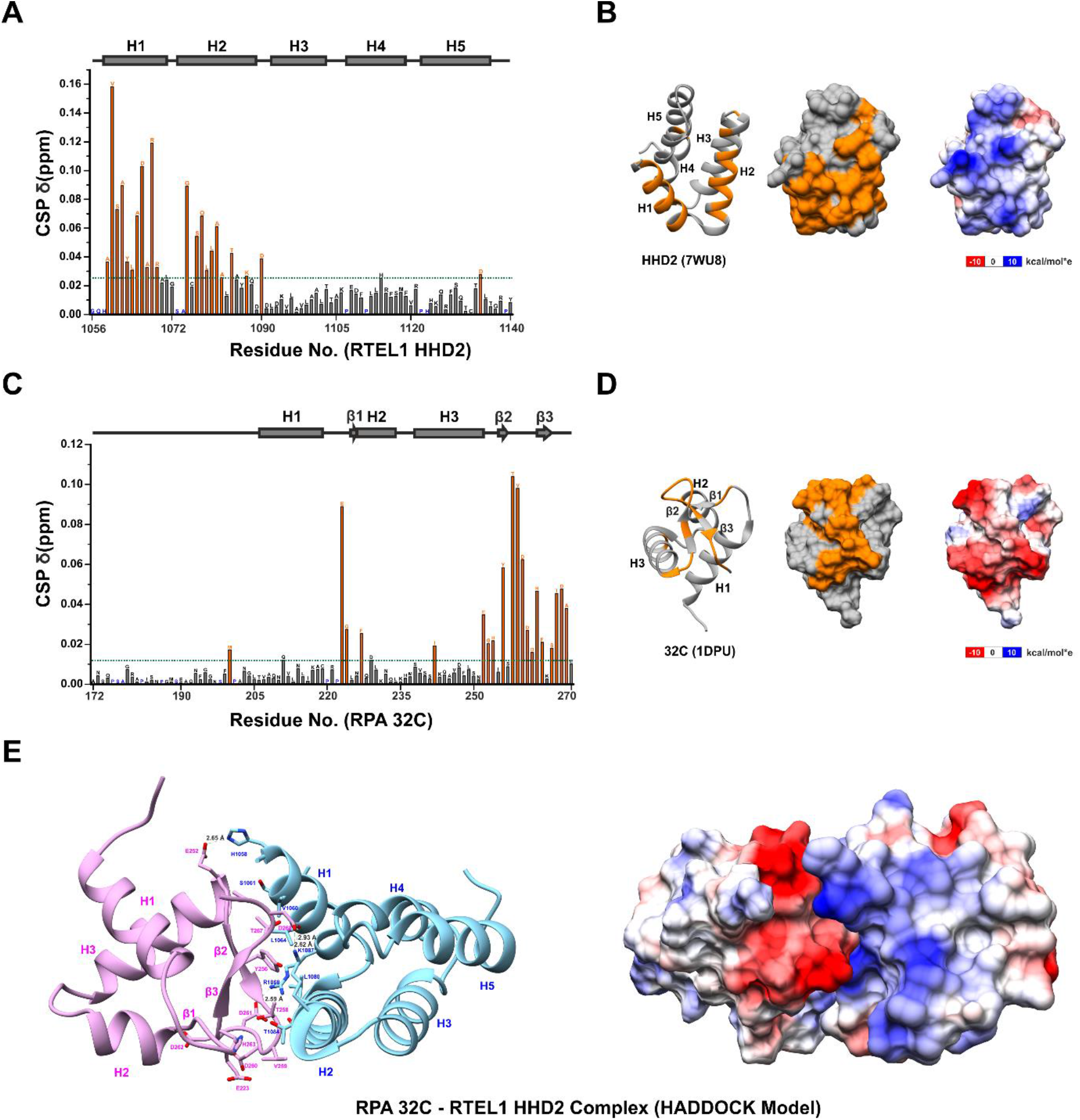
The binding interface of RTEL1-RPA interaction. **(A)** Quantification of CSPs in HHD2 upon titration with RPA 32C (at 1:4 molar ratio). Residues with more than average CSP (green dash line) are marked as an orange bar and considered as significantly perturbed residues. Proline and unassigned residues are marked in blue. The secondary structure corresponding to HHD2 is shown at the top. **(B)** Significantly perturbed residues (orange) are marked on the structure of HHD2 (ribbon - left and surface - middle forms). Most of these residues lie on the positively charged surface (right) of HHD2. **(C)** Quantification of CSPs in 32C upon titration with HHD2 (at 1:4 molar ratio). Residues with more than average CSP (green dash line) are marked as an orange bar and considered as significantly perturbed residues. Proline and unassigned residues are marked in blue. The secondary structure corresponding to 32C is shown at the top. **(D)** Significantly perturbed residues (orange) are marked on the structure of 32C (ribbon - left and surface - middle forms). Most of these residues lie on the negatively charged surface (right) of 32C. **(E)** HADDOCK model of RTEL1 HHD2 – RPA 32C complex with indicated interface residues. Positively (blue) and negatively (red) charged interacting surfaces of HHD2 and 32C, respectively, are shown.

Similarly, the observed CSPs in the titration of ^15^N-labelled 32C with HHD2 (Figure 3C) were calculated and plotted (Figure 5C) to map the HHD2 binding surface on the RPA 32C domain. A previously determined structure (PDB ID 1DPU) and deposited chemical shifts (BMRB entry 4460) of RPA 32C were used for the analysis (Mer et al., 2000). The residues (E223, G224, F227, I242, E252, G253, H254, Y256, T258, V259, D260, D261, D262, H263, F264, S266, T267, D268, and A269) that showed perturbation more than the average CSPs upon HHD2 binding were mainly present in the C-terminal β-sheet and the adjacent loops of 32C (Figure 5D). Interestingly, this region of 32C has been shown to interact with several proteins involved in the DNA repair and replication processes (Figure S3A and S3B) (Dueva and Iliakis, 2020). Based on these observations, we conclude that the RTEL1–RPA interaction is mediated through the conserved binding surfaces on HHD2 of RTEL1 and winged-helix domain 32C of RPA.

Electrostatic surface potential analysis revealed that the positively charged surface of HHD2 (Figure 5B) interacts with the negatively charged surface of 32C (Figure 5D). To get molecular insight into this interaction, we generated the NMR CSPs data-driven HADDOCK model of the HHD2–32C complex (Figure 5E and Supplementary Table S1). The largest cluster (cluster 1) contains 79 models, out of the 153 final clustered models, with the best HADDOCK score (−78.6 ± 3.0) and Z-score (−1.9), suggesting that the selected HADDOCK cluster is of good quality. Therefore, we chose this cluster to represent the HHD2–32C complex. A closer inspection of the selected model showed that hydrophobic and ionic interactions stabilize the HHD2–32C complex (Figure 5E). Upon complex formation, the total buried surface area is 1322.7 ± 29.4 Å^2^ (Supplementary Table S1).

### HHD2 of RTEL1 interacts with DNA

The HRDC domain in WRN and BLM DNA helicases regulates its helicase activity through binding with DNA (Estep and Brosh, 2018; Kim and Choi, 2010). We hypothesize that HHDs of RTEL1 may have similar functions and can potentially interact with DNA.

We performed NMR titrations of individual HHDs using a 22-mer ssDNA (ssDNA-22) and a 22 base-paired double-stranded DNA (dsDNA-22). There were no significant CSPs in the ^1^H-^15^N HSQC spectra of HHD1 upon ssDNA titrations (Figure S4A). In the case of dsDNA titrations, we observed minor perturbations in chemical shifts of a few residues of HHD1 (Figure S4B). These results suggest that HHD1 of RTEL1 either does not bind ssDNA (ssDNA-22) or binds dsDNA weakly (dsDNA-22).

Interestingly, in the case of the titrations of HHD2 with DNA, we observed distinct CSPs upon the addition of both ssDNA and dsDNA. HHD2 residues that showed large CSPs upon ssDNA (Figure 6A and 6B) and dsDNA binding (Figure S5A and S5B) were mapped to the helices H1, H2, and H4 of the HHD2 structure (Figure 6C and 6D; and Figure S5C and S5D).

**Figure 6.**
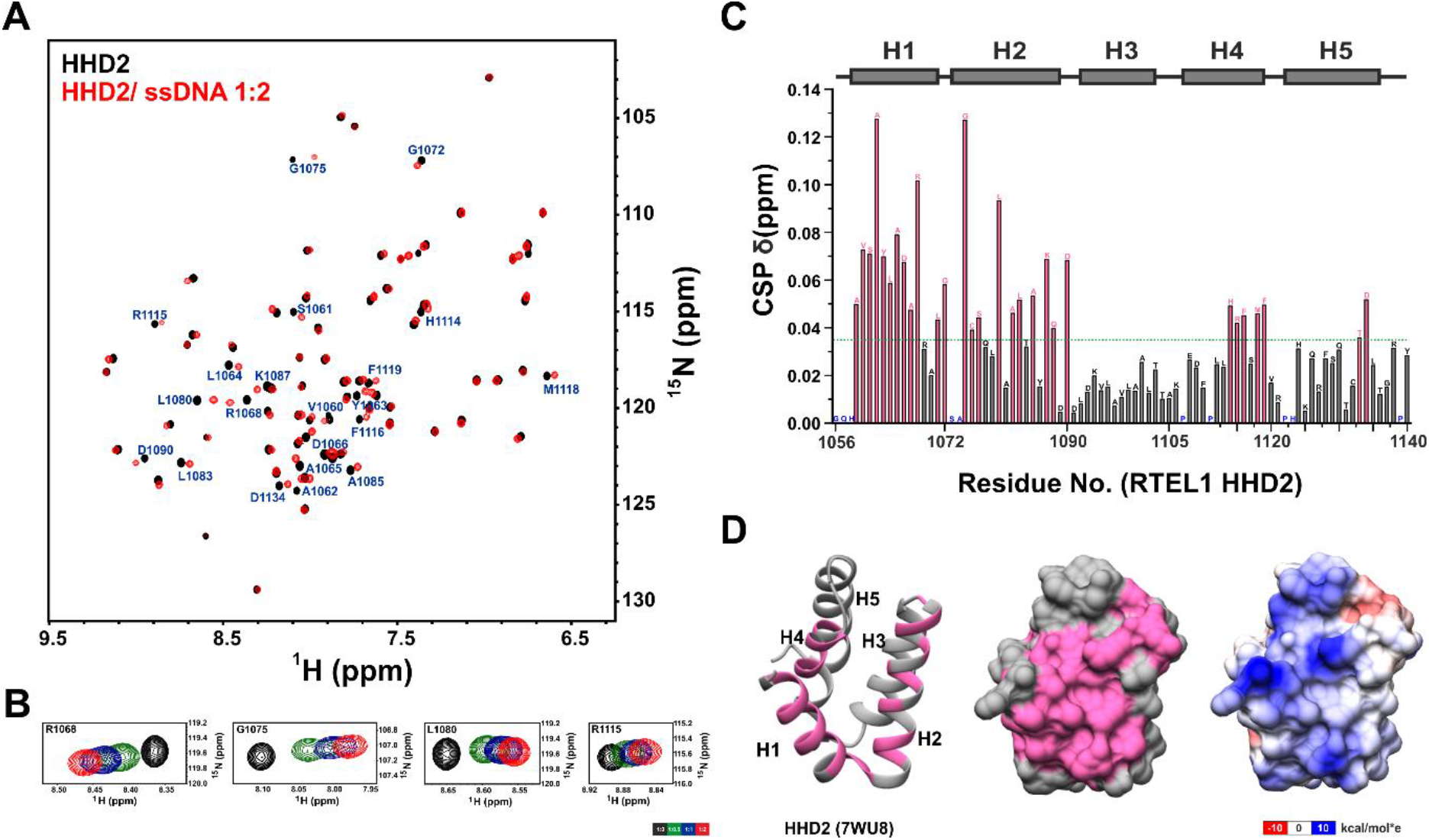
NMR titration of HHD2 and ssDNA. **(A)** Overlay of ^1^H -^15^N HSQC spectra of HHD2 in the absence (black) and presence (red) of ssDNA-22 (at 1:2 molar ratio). Residues that showed large CSPs are labeled. **(B)** ^1^H-^15^N cross-peaks trajectory of representative residues R1068, G1075, L1080, and R1115 of HHD2 upon titration with ssDNA-22 at indicated molar ratios. **(C)** Quantification of CSPs in HHD2 upon titration with ssDNA-22 (at 1:2 molar ratio). Residues that showed more than average CSP (green dash line) are marked as a pink bar and considered as significantly perturbed residues. Proline and unassigned residues are marked in blue. The secondary structure of HHD2 is shown at the top. **(D)** Significantly perturbed residues (pink) are marked on the structure of HHD2 (ribbon – left and surface – middle form). Most of these residues lie on the positively charged surface (right) of HHD2.

To further understand the HHD2–DNA interaction, we performed isothermal titration calorimetry (ITC) experiments. ITC experiments allow the determination of thermodynamic parameters such as a change in enthalpy (ΔH), change in entropy (ΔS), equilibrium dissociation constant (K_d_), and stoichiometry (n) of interaction under a set of experimental conditions. The ITC results (Figure 7A-7C) revealed that HHD2 interacts with both ssDNA and dsDNA in enthalpically driven binding with dissociation constants (K_d_) of 6.49±0.32 μM and 4.67±0.32 μM, respectively (Table 2). These results unequivocally showed that HHD2 of RTEL1 interacts with DNA. This is the first report of an HHD binding to DNA.

**Table 2.**
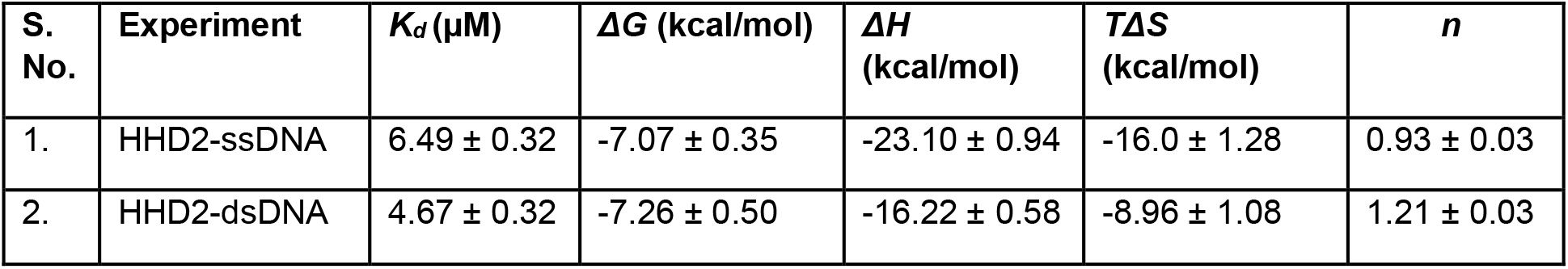
Equilibrium dissociation constants (K_d_s) and other thermodynamic parameters for RTEL1 HHD2 and DNA interactions using ITC experiments.

**Figure 7.**
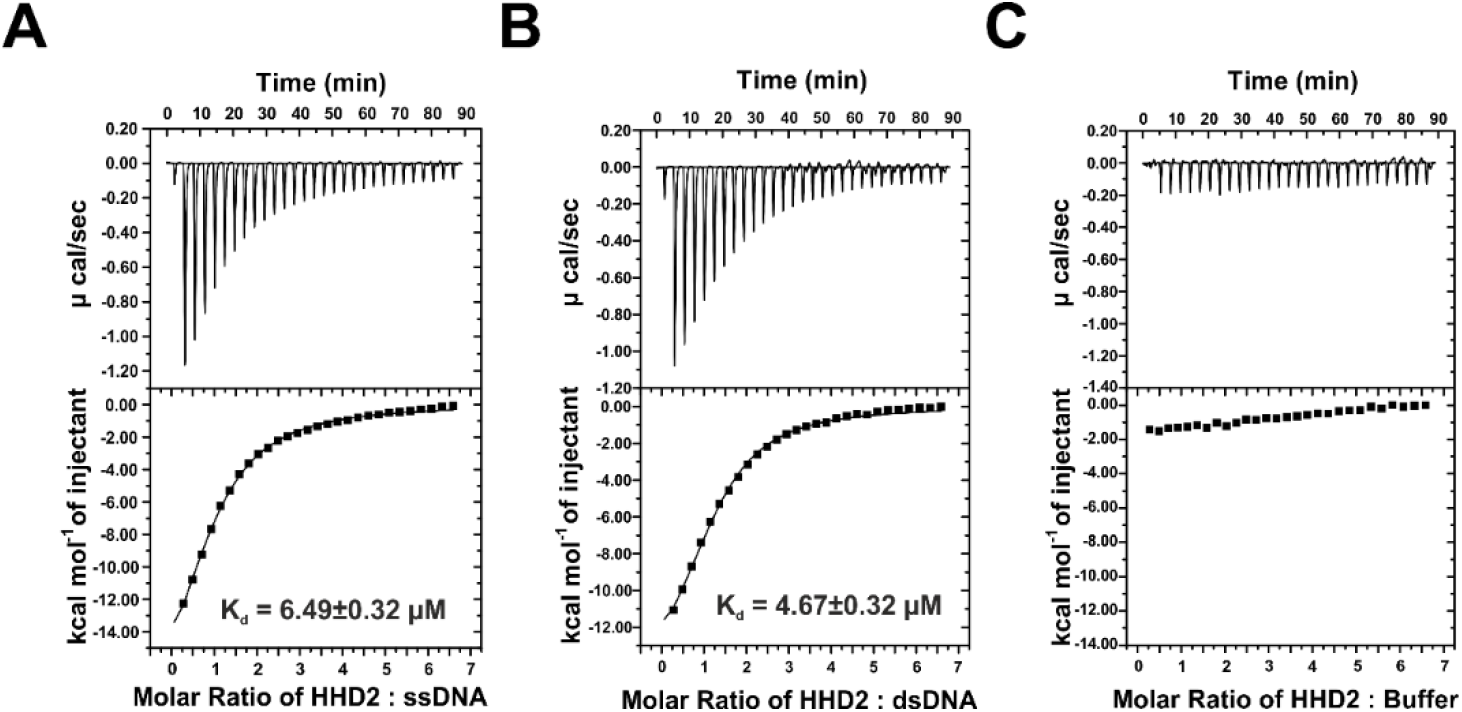
Isothermal titration calorimetry of HHD2 and DNA. **(A)** Raw and fitted ITC binding isotherms for the interaction of HHD2 with ssDNA-22. The equilibrium K^d^ obtained upon fitting the raw data is mentioned. **(B)** Raw and fitted ITC binding isotherms for the interaction of HHD2 with dsDNA-22. The equilibrium K^d^ obtained upon fitting the raw data is mentioned. **(C)** Raw and fitted ITC binding isotherms for the control titration of HHD2 with ITC buffer.

### RPA 32C and DNA compete for the same binding site on RTEL1 HHD2

We observed that the RPA 32C binding surface overlaps with the DNA binding surface in the HHD2 structure (Figure 8A). RPA 32C binds mainly to the solvent-exposed surface of helix H1 and H2, while DNA binds in the pocket formed by helix H1, H2, and H4, suggesting HHD2 adapts to bind DNA and 32C on the same surface. We hypothesized that RPA 32C and DNA may bind competitively to HHD2 of RTEL1. We performed an NMR-based competitive experiment to address this hypothesis and recorded a ^1^H-^15^N HSQC spectrum of ^15^N-labeled HHD2 with 32C. The HHD2–32C complex was then titrated with the ssDNA-22. We visualized this competitive binding by monitoring the resonance cross peak of three representative residues R1068 (binds both 32C and DNA), A1081 (binds only 32C), and F1119 (binds only DNA) (Figure 8A). The addition of DNA resulted in resonance peaks coming to the position of the HHD2–DNA complex, thus suggesting the competitive displacement of 32C by the DNA (Figures 8B and 8C).

**Figure 8.**
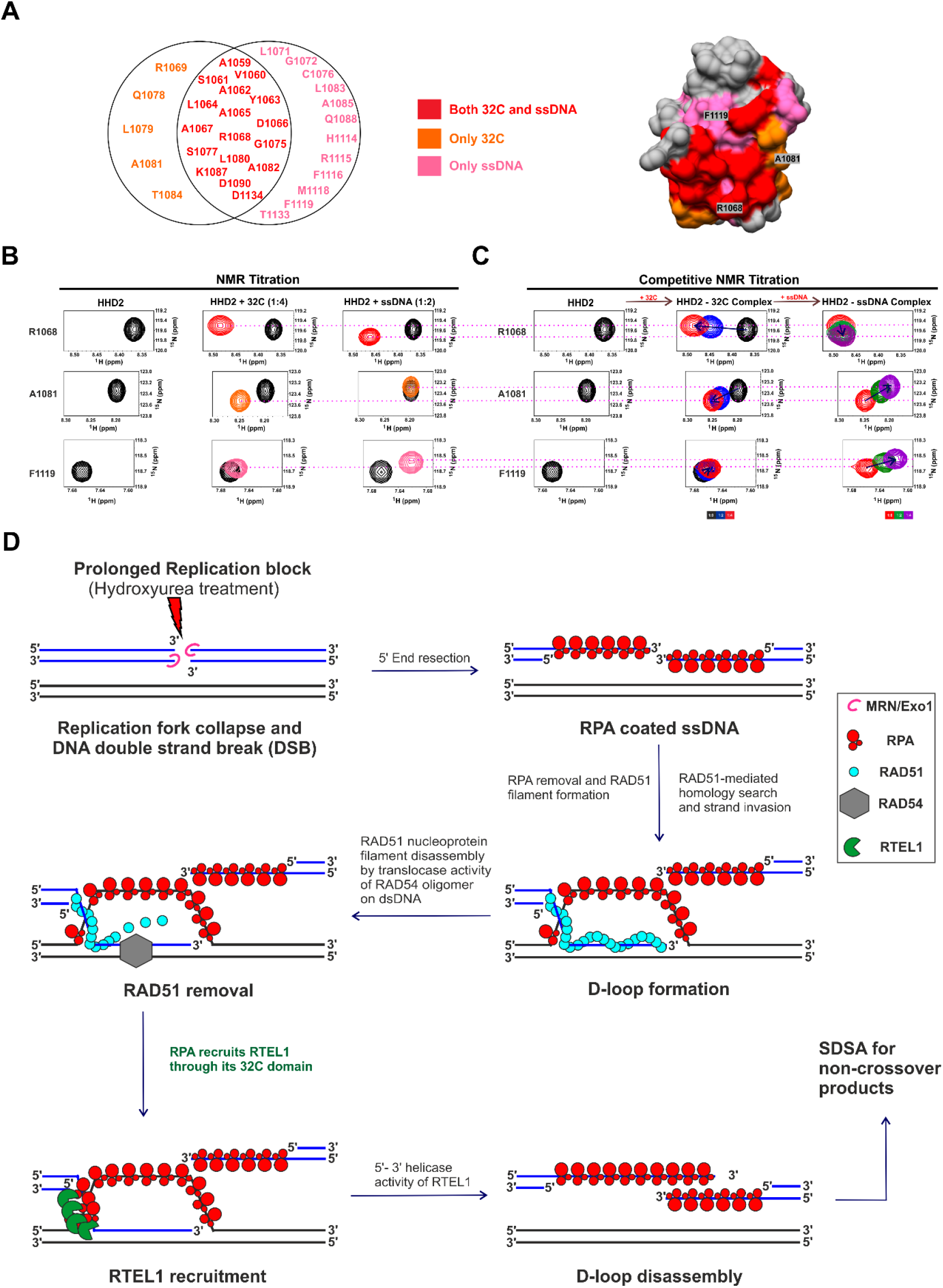
The interplay among RTEL1, RPA, and DNA. **(A)** A subset of significantly perturbed residues of HHD2 upon binding to only RPA 32C (orange), only ssDNA (pink), and both 32C and ssDNA (red) are shown. One representative residue from each subset is denoted on the surface-filled structure of HHD2. **(B)** ^1^H-^15^N HSQC cross-peaks trajectory of representative residues R1068, A1081, and F1119 of HHD2 upon individual titration with RPA 32C and ssDNA-22 at indicated molar ratios. **(C)** Competitive NMR titration of ^15^N-HHD2 with RPA 32C and ssDNA. Arrow indicates the direction of movement of the cross-peaks upon titration at indicated molar ratios. NMR titration of the HHD2–32C complex with DNA showed that the cross-peak of A1081 comes back to the free HHD2/HHD2-DNA complex position while the cross-peaks of F1119 and R1068 follow the path towards the HHD2-DNA complex. **(D)** Proposed model for RPA-mediated recruitment of RTEL1 to perform efficient D-loop disassembly. Prolonged replication block by hydroxyurea treatment eventually leads to DNA double-strand break (DSB). MRN/Exo1 (pink) complex executes 5’ end resection to generate the 3’ overhang, which quickly gets coated by RPA (red). RAD51 (cyan) nucleoprotein filament invades the homologous dsDNA to form D (displacement)-loop. RAD54 (grey) translocase activity helps in the removal of RAD51. RPA, present on the displaced ssDNA and invading strand, recruits the RTEL1 (green) on D-loop through its 32C domain. The 5’-3’ helicase activity of RTEL1 helps disassemble the D-loop. It thus directs the recombination intermediate to follow the synthesis-dependent strand-annealing (SDSA) pathway of homologous recombination to generate the non-crossover products.

## DISCUSSION

### RTEL1 performs DNA damage repair through RPA-mediated recruitment at D-loop

RTEL1 acts as an anti-recombinase by dissociating the D-loop structure during the homologous recombination (Barber et al., 2008; Youds et al., 2010). Although a model describing the role of RTEL1 in the synthesis-dependent strand annealing (SDSA) pathway of homologous recombination was proposed (Uringa et al., 2011), how RTEL1 is recruited at the D-loop had remained unknown.

Here, we show the nuclear co-localization of RTEL1 and RPA under DNA damage conditions. Co-immunoprecipitation experiments showed that RPA and RTEL1 interact. We observed that deletion of the HHD1+2 region from RTEL1 significantly reduced RPA–RTEL1 interaction, however, it didn’t abolish the interaction with RPA completely (Figure 2C). Since both RPA and RTEL1 are multi-domain proteins, we postulate that there might be the possibility of other interaction sites in addition to the HHD2 binding.

Several RPA interacting proteins like RAD52, XPA, SV40 Tag, WRN, etc., show multivalent binding with RPA; primary interaction through RPA 70N or RPA 32C and secondary interaction within the tandem RPA 70A-70B domain (Caldwell and Spies, 2020; Sugitani and Chazin, 2015; Yeom et al., 2019). Using extensive NMR titration-based experiments, we probed the binding of HHD1 and HHD2 of RTEL1 with different domains of RPA. The results showed that the RTEL1 HHD2 interacts exclusively with the RPA 32C domain (Figure S2A-S2F and Figure 3A-3D).

Cellular signalling and stress response pathways often involve weak (K_d_ in mM to μM range) and transient protein-protein interactions (Cohen and Pielak, 2017; Sugitani and Chazin, 2015; Sukenik et al., 2017). RPA is the first responder to ssDNA and acts as a hub protein to recruit multiple specific factors (helicases, translocase, nuclease, etc.) to regulate DNA replication, repair, and recombination processes (Chen and Wold, 2014). The RPA-mediated protein-protein interactions are weak (μM affinity) and transient in nature (Sugitani and Chazin, 2015; Yeom et al., 2019), which is reflected in the fast-exchange kinetics of NMR titration between RTEL1 HHD2 and RPA 32C (Figure 3B and 3D).

RPA binds ssDNA with high affinity (K_d_ in nM) owing to the presence of multiple <u>D</u>NA <u>b</u>inding OB-fold <u>d</u>omains (DBDs) (Wold, 1997). Microscopic association and dissociation of individual DBDs lead to conformational dynamics of the ssDNA-RPA complex. The RPA interacting proteins selectively modulate the conformational dynamics of individual DBDs (Pokhrel et al., 2019). Thus, RPA can pass the ssDNA to the interacting partner that has lower binding affinity for DNA (e.g., ∼ 6 μM K_d_ in case of RTEL1 HHD2 – DNA binding).

Based on the existing literature and our results presented here, we proposed a model describing the role of RPA in the recruitment of RTEL1 at the D-loop (Figure 8D). RPA is present on the displaced ssDNA of the D-loop structure at repair sites (Eggler et al., 2002; Li and Heyer, 2008). Once the RAD54, HELQ-1, and RFS-1 displace the RAD51 from the invading strand of the D-loop (Krejci et al., 2012; Solinger and Heyer, 2001; Ward et al., 2010), transiently exposed ssDNA of the invading strand gets occluded by RPA. The RPA, present on the invading strand and the displaced strand of the D-loop, is ready to recruit specific downstream factors. The RPA 32C and RTEL1 HHD2 interaction will recruit RTEL1 at the ss-dsDNA junction of the invading strand of the D-loop. A conformational change in RPA (Pokhrel et al., 2019) upon interaction with RTEL1 will expose the ssDNA, making it available for RTEL1 binding. It is noteworthy that the RTEL1 HHD2 domain has a high affinity for ssDNA compared to its affinity for the RPA 32C (Figure 8C), which will facilitate the RPA-mediated recruitment of RTEL1 at the D-loop. Upon recruitment, RTEL1 unwinds the DNA at ss-dsDNA junction using its 5’-3’ DNA helicase activity and thus help resolve the D-loop structure (Figure 8D).

In an in vitro assay, RTEL1 was shown to preferentially dissociate the D-loops with 3’ invasion. Interestingly, the efficient unwinding required the presence of the RPA (Youds et al., 2010). Our results and the proposed model of RPA-mediated recruitment of the RTEL1 helicase at the D-loop structure could explain the mechanism behind this observation.

### Functional plasticity of Harmonin-homology domains

HHDs belong to a newly classified group of αα-hub domains found in several proteins (Colcombet-Cazenave et al., 2021). The αα-hub domains typically consist of 3-5 α-helices. Hub proteins mediate fidelity in signaling and larger protein complexes. HHDs consist of five α-helices proposed to act mainly as a protein-protein interaction domain. The HHDs are associated with the PDZ domain in Harmonin, Whirlin, Delphilin, and PDZD7, with the PID domain in CCM2, and with the helicase and RING domains in RTEL1 (Figure S6A), suggesting that HHDs have evolved to provide context-dependent functions in different proteins (Bugge et al., 2021; Colcombet-Cazenave et al., 2021; Staby et al., 2021).

Although the structure of HHDs in these proteins bears high structural similarity (Figure S6B) the functions of these proteins are diverse and entirely unrelated (Colcombet-Cazenave et al., 2021). The Harmonin and CCM2 HHDs interact with partner proteins (Colcombet-Cazenave et al., 2021). The mouse PDZD7 HHD-region interacts with lipids (Lin et al., 2021). Here, we report the DNA and protein binding properties of the HHD2 domain of RTEL1. Therefore, HHDs display remarkable functional plasticity to interact with diverse biomolecules (i.e., protein, lipids, and DNA).

### Unique surface charge distribution imparts dual function to RTEL1 HHD2

The X-ray crystal structure of the HHD2 domain of RTEL1 reported here is the first structure of any domains of RTEL1. HHD1 and HHD2 domains of RTEL1 fold into a common globular bundle of five helices (Figure S6B). However, the amino acids on the domain surfaces are poorly conserved, yielding distinct surface properties. The isoelectric point (pI) of HHD1 and HHD2 is 6.31 and 8.73, respectively (Figure S6C). HHD1 has an acidic surface and a largely neutral opposite surface. However, HHD2 consists of distinct basic and acidic surfaces (Figure S6D), suggesting that these two tandem domains may have distinct roles in RTEL1.

The tandem HHDs of RTEL1 have the same fold, but specific surface properties suggesting its evolutionary significance. In the case of plants and invertebrates, RTEL1 has only one HHD, while vertebrates have two tandem HHDs in their RTEL1 protein (Colcombet-Cazenave et al., 2021; Faure et al., 2014). We have shown that the tandem HHD1 and HHD2 domains are independently folded and separated by a 75 residue-long disordered linker regions. The distinct surface property and spatial separation through unstructured and long linker regions may be advantageous for their interaction with different proteins to coordinate multiple cellular functions. For example, only HHD1 was shown to interact with SLX4 (Takedachi et al., 2020). A short alpha-helical C-terminal extension of HHD1 is critical for interaction with SLX4, while the HHD2 has no such extension. The distinct positively charged surface of HHD2 helps mediates its interaction with the negatively charged surface of the RPA 32C and DNA.

The HHD2 interacts with the 32C domain of RPA and DNA using a common surface (helices H1 and H2). Interestingly, equivalent helices in the Harmonin and CCM2 HHDs were reported to constitute the protein-binding sites in them (Colcombet-Cazenave et al., 2021). Like several RPA interacting proteins, the HHD2 of RTEL1 interacts with the conserved binding surface of RPA 32C (Figure S3A and S3B). Therefore, we conclude that the HHD2 of RTEL1 is a unique hub domain capable of mediating protein-protein and protein-DNA interactions.

Interestingly, the HRDC domain of RecQ DNA helicases is also all-helical (consisting of five α-helices and one 3_10_-helix) like the HHDs of the RTEL1 (Kim and Choi, 2010). The HRDC domains interact with DNA (Kim and Choi, 2010), facilitating the recruitment of these DNA helicases to the site of DNA repair (Croteau et al., 2014). The HRDC–DNA interaction confers specificity and modulates the G-quadruplex unwinding activity of REQ helicase in *E. coli* (Teng et al., 2020). Based on the results from this study, we suggest that the HHDs could provide the HRDC equivalent roles in RTEL1.

### Future perspective of the interplay among RTEL1, RPA, and DNA

Since RTEL1 HHD2 interacts with both ssDNA and dsDNA with similar affinity (Table 2), we postulate that HHD2 has the potential to bind at the ss-dsDNA junction formed during DNA metabolism processes in the cell. We have shown the competitive binding of RPA 32C and DNA on the HHD2 of RTEL1 (Figure 8C). This interplay of RPA-RTEL1-DNA interactions will have implications for understanding the mechanism of RTEL1 in the DNA replication, repair, and recombination processes. RTEL1 interacts with the PCNA through its PIP-box motif and helps genome-wide replication (Vannier et al., 2013). RTEL1 facilitates bypass of the DNA-protein cross-links (DPCs) by replicative helicase CMG; interestingly, PIP box of the RTEL1 is not required for this function (Sparks et al., 2019). This suggests additional mechanisms (apart from the PCNA-mediated) of RTEL1 recruitment at the replication fork. Future studies may unravel the possibility of RPA-mediated recruitment of the RTEL1 at the replication fork. It would be interesting to explore the effect of the abolishment of the RTEL1-RPA interaction on DNA repair, genome-wide DNA replication, DPC bypass, and telomere maintenance.

## METHODS

### Plasmid construction

DNA sequences corresponding to HHD1 (residues 886-978) and HHD2 (residues 1056-1140) were cloned in the pET-28a vector as described earlier (Kumar et al., 2022). The DNA sequences encoding the tandem harmonin homology domains (HHD1+2) (residues 886– 1140) of human RTEL1 (Uniprot identifier Q9NZ71-6) were PCR amplified using a human RTEL1 cDNA clone as a template and subcloned into *E. coli* expression vector pET-28a (+) between NdeI and XhoI restriction endonuclease sites.

The DNA sequences encoding the 70N (residues 1–120 of RPA1), 70A (residues 181–291 of RPA1), 70B (residues 299–422 of RPA1), and 32C (residues 172–270 of RPA2) domains of human RPA were PCR amplified using p11d-tRPA plasmid (Addgene plasmid102613) (Henricksen et al., 1994) as a template and subcloned into *E. coli* expression vector pET-15b between NdeI and BamHI restriction endonuclease sites.

The forward and reverse primer sets for each construct are listed in the supplementary Table S2. Phusion High-Fidelity DNA polymerase (New England Biolabs; Cat. No. M0530) was used for PCR amplification of each construct. The recombinant plasmids were amplified in *E. coli* DH5α cells and isolated through the QIAprep Spin Miniprep Kit (Qiagen; Cat. No. 27106). All the cloned plasmids (Supplementary Table S3) encode a hexa-histidine (6xHis) purification tag at the N-terminus of the protein sequence.

### Protein expression and purification

#### RTEL1 HHD1, HHD2, and HHD1+2

RTEL1 HHD1 and HHD2 domains were expressed and purified as described previously (Kumar et al., 2022). In summary, the following protocol was followed for the purification of HHD1, HHD2, and HHD1+2. Plasmids containing HHD1, HHD2, and HHD1+2 were individually transformed into *E. coli* Rosetta (DE3) cells. Cells were grown in either the LB (for unlabeled protein) or M9 minimal media (for uniformly ^15^N-labeled protein for NMR) in the presence of Kanamycin and Chloramphenicol antibiotics. ^15^N NH_4_Cl (Cambridge Isotope Laboratories; Cat. No. NLM-467-10) was used (1g/L of the media) as a sole source of nitrogen in case of minimal media preparation. Protein expression was induced at OD_600_ of 0.8-1 by adding 1 mM of IPTG, and the culture was incubated at 20°C for 18 h. Cells were harvested and lysed using sonication under lysis buffer (50 mM Tris pH 8 at 4°C, 500 mM NaCl, 10 % Glycerol, 0.02 % NaN_3_). EDTA-free Protease Inhibitor Cocktail tablets (Roche; Cat. No.11836170001) and 1 mM PMSF (Roche; Cat. No.10837091001) were added before lysing the resuspended cells through sonication. The lysate was subjected to centrifugation at 13,000 rpm for 60 min at 4°C, and the supernatant was collected. 0.1% (v/v) polyethyleneimine (PEI) precipitation followed by another round of centrifugation was carried out to remove any nucleic acid contamination in the case of HHD2 and HHD1+2.

For Ni^2+^–NTA affinity chromatography, the supernatant was loaded on the His Trap FF column (GE) pre-equilibrated with lysis buffer. Extensive column wash was performed with wash buffer (50 mM Tris pH 8 at 4°C, 500 mM NaCl, 20 mM Imidazole, 10 % Glycerol, 0.02 % NaN_3_) to remove the non-specifically bound proteins. 6xHis-tagged proteins were eluted with imidazole in the elution buffer (50 mM Tris pH 8 at 4°C, 500 mM NaCl, 250 mM Imidazole, 10 % Glycerol, 0.02 % NaN_3_).

For performing the ion-exchange chromatography, Ni^2+^–NTA eluted proteins were subjected to buffer exchange in buffer A (20 mM Tris pH 7.4 at 4°C, 50 mM NaCl, 2 mM DTT, 5% glycerol, 0.02% NaN_3_) and loaded on the ion-exchange column (HiTrap Q or HiTrap Heparin column, GE) pre-equilibrated with buffer A. The bound protein was eluted with a linear gradient of 1M NaCl containing buffer B (20 mM Tris pH 7.4 at 4°C, 1 M NaCl, 2 mM DTT, 5% glycerol, 0.02% NaN_3_).

Finally, size-exclusion chromatography (SEC) was carried out on a Superdex 75 column (HiLoad 16/600, prep grade; GE) pre-equilibrated with the NMR buffer (20 mM Tris-HCl pH 7.4 at 25°C, 50 mM NaCl, 2 mM DTT, 0.02% NaN_3_), or crystallization buffer (20 mM Tris-HCl pH 7.4 at 25°C, 250 mM NaCl, 5% Glycerol, 2 mM DTT, 0.02% NaN_3_), or ITC buffer (10 mM Potassium phosphate pH 6.5 at 25°C, 50 mM KCl, 0.02% NaN_3_) depending on the intended use of the protein for the subsequent experiments.

SDS-PAGE analysis was performed to ascertain the purity, molecular weight (relative to Precision Plus Protein Standards Dual colour; Bio-Rad, Cat. No.161-0394), and integrity of the purified proteins (Figure S1A). Proteins were concentrated using a centrifugal filter unit (3 kDa MWCO, Merck Millipore; Cat. No. UFC900324) at 3500 rpm and 4°C. The HHD2 samples were concentrated at 15°C, owing to its less solubility at 4°C. The concentrations of the protein were calculated by ultraviolet (UV) absorbance at 280 nm using a UV spectrophotometer (Eppendorf), and the calculated molar extinction coefficients of HHD1, HHD2, and HHD1+2.

#### Full-length RPA trimeric complex

RPA complex was expressed and purified as per the protocol from the M.S. Wold lab (Binz et al., 2006; Henricksen et al., 1994). p11d-tRPA (123) plasmid (Supplementary Table S3) was transformed into *E. coli* BL21(DE3) strain. Cells were grown in the presence of Ampicillin antibiotics. Protein expression was induced at OD_600_ of 0.7-0.8 by adding 0.3 mM of IPTG, and the culture was incubated at 37°C for 3 h. Protein purification was performed through Affi-gel blue (Bio-Rad; Cat. No.153-7302), Hydroxyapatite (Bio-Rad; Cat. No.130-0420), and Q-column (GE). 1.5 M NaSCN (Sigma; Cat. No. 251410), 80 mM Potassium phosphate, and 300 mM KCl buffer were used to elute the RPA (Figure S1D) from Affi-Gel blue, Hydroxyapatite, and Q-column, respectively. Finally, dialysis of RPA was carried out in the NMR buffer containing 100 mM NaCl (20 mM Tris-HCl pH 7.4 at 25°C, 100 mM NaCl, 2 mM DTT, 0.02% NaN_3_).

#### RPA 70N, 70A, 70B, and 32C

RPA 70N, 70A, 70B, and 32C domains were expressed and purified as described previously (Bhattacharya et al., 2004; Bochkareva et al., 2005; Lee and Park, 2016; Mer et al., 2000) with a few modifications. Plasmids carrying coding DNA for different RPA domains were transformed into *E. coli* BL21 (DE3) (for 70N domain) or BL21 (DE3) pLysS strains. Cells were grown in either the LB (for unlabeled protein) or M9 minimal media (for uniformly ^15^N labeled protein for NMR) in the presence of Ampicillin / Ampicillin and Chloramphenicol antibiotics. ^15^N NH_4_Cl (1g/L of the media) was used as a sole source of nitrogen in case of minimal media preparation. Protein expression was induced at OD_600_ of 0.8-1 by adding 1 mM of IPTG, and the culture was incubated at 37°C for 4 h (70N incubated at 20°C for 18 h). The first step of purification involved His-tag - Ni^2+^-NTA affinity chromatography for all the domains. In the next step, RPA 70A and 70B were purified using cation exchange (Heparin column). RPA 32C was purified using anion exchange (Q column). All the domains were purified using SEC on a Superdex 75 column at the final step of purification and eluted in the NMR buffer (20 mM Tris-HCl pH 7.4 at 25°C, 50 mM NaCl, 2 mM DTT, 0.02% NaN_3_).

SDS-PAGE analysis was performed to ascertain the purity, molecular weight (relative to protein standard), and integrity of the purified proteins (Figure S1D). Proteins were concentrated using a centrifugal filter unit (3 kDa MWCO, Merck Millipore) at 3500 rpm and 4°C. The concentrations of the protein were calculated by ultraviolet (UV) absorbance at 280 nm using a UV spectrophotometer (Eppendorf), and the calculated molar extinction coefficients of 70N, 70A, 70B, and 32C.

### Co-immunoprecipitation (Co-IP)

#### Cell culture and transfection

HEK293T cells were grown in low glucose DMEM supplemented with 10% foetal bovine serum and an antibiotic-antimycotic mix. Cells were maintained at 37°C and 5% CO_2_ in a humidified incubator. For transfection, cells were grown to 70–80% confluence, and the plasmids (pcDNA3.1(+)-N-HA empty vector, pcDNA3.1(+)-N-HA-RTEL1, and pcDNA3.1(+)-N-HA-RTEL1-ΔHHD1+2) (Supplementary Table S3) were transfected using Lipofectamine® 2000 reagent (Invitrogen; Cat. No.<u>11668019</u>) according to the manufacturer’s protocol. 48 h post-transfection, the cells were harvested.

#### Preparation of cell lysates

Cells were harvested from culture dishes using a cell scraper and lysed with ice-cold cell lysis buffer (20 mM Tris–HCl pH 7.5, 100 mM NaCl, 1 mM EDTA, 1 mM EGTA, 1% Triton X-100, 2.5 mM sodium pyrophosphate, 1 mM sodium orthovanadate, 1 mM PMSF and 1X protease inhibitor cocktail). The lysates were cleared by centrifugation at 12,000 rpm for 10 min at 4°C, and the supernatant was collected in a fresh tube.

#### Immunoprecipitation, electrophoresis, and immunoblotting

Protein lysates were quantified using the Bradford assay, and 500-750 μg lysate was pre-cleared with Protein G agarose beads (Roche; Cat. No.11719416001) for 1 hour at 4°C with rotation at 5 rpm in a rotator. Pre-cleared lysates were incubated with 3 μg of specific antibody overnight at 4°C with rotation at 5 rpm in a rotator. The immunoprecipitated complexes were captured on protein G agarose beads. After brief centrifugation, the supernatant was discarded, and beads were washed thrice with ice-cold PBS (1X) at 500 rpm and 4°C for 30 sec. After boiling the beads at 95°C for 5 min in 2X Laemmli sample buffer with 5% β-mercaptoethanol (Bio-Rad), the immunoprecipitated protein was resolved on 10% SDS-PAGE gels and then transferred onto 0.45 μm PVDF membrane (GE) by overnight wet transfer. The blot was blocked with 5% skimmed milk solution (prepared in TBST, Tris-buffered saline supplemented with 0.05% Tween 20) for 1 h at room temperature, followed by incubation at 4°C overnight with a specific primary antibody prepared in 5% BSA or skimmed milk in TBST (dilution range: 1:500 to 1:1,000). Bound antibodies on the blot were probed with HRP-conjugated secondary antibody at RT for 1 h. Between each step, the blots were washed thrice for three minutes each with TBST. Chemiluminescence was detected using ECL western blotting substrate (Bio-Rad), and the images were acquired using a chemiluminescence imager (Bio-Rad).

### Immunofluorescence microscopy

HeLa cells were grown on sterilized glass coverslips placed inside a 12-well plate. 4 mM Hydroxyurea (Sigma; Cat. No. H8627) was added to the 60% to 80% confluent cultures for 48 h before harvesting. The cells were fixed using 4% paraformaldehyde for 10–15 min at room temperature. Permeabilization was carried out with 0.2% Triton X-100 for 5 min at room temperature, followed by incubation with blocking buffer (5% BSA in PBS containing 0.1% Tween 20) for 60 min. The cells were incubated overnight at 4°C with indicated primary antibodies (prepared in the blocking buffer at dilutions ranging from 1:50–1:150) followed by incubation with species-specific Alexa Fluor-conjugated secondary antibodies for 1 h at room temperature. The nuclei were stained with Hoechst 33342 (Cayman chemical; Cat. No. 15547) by incubation for 15 min at room temperature. The cells were washed twice with 1X PBS between each step. Finally, the cells were mounted on a clean glass cover slide using the Fluoromount-G Aqueous Mounting Medium, and the images were acquired using Zeiss LSM 880 confocal microscope. The images were further processed and analyzed with ZEN 3.5 (Blue edition).

### Antibodies

Anti-RTEL1, produced in Rabbit (Sigma; Cat. No. HPA067329); Anti-RPA1, produced in Mouse (Sigma; Cat. No. SAB1406399); Anti-RPA 32 kDa subunit (9H8), produced in Mouse (Santa Cruz Biotechnology; Cat. No. sc-56770); Anti-HA tag, produced in Rabbit (Abcam; Cat. No. ab9110); Anti-beta Actin, produced in Mouse (Abcam; Cat. No. ab8226); Anti-Rabbit IgG, HRP-conjugate (Millipore; Cat. No. 12-348); Anti-Mouse IgG, HRP-conjugate (Cell Signalling Technology; Cat. No. 7076); Anti-Rabbit IgG, Alexa Fluor™ 488-conjugate (Invitrogen; Cat. No. A-11008); Anti-Mouse IgG, Alexa Fluor™ 594-conjugate (Invitrogen; Cat. No. A-11032)

### Crystallization and X-ray structure determination

The SEC fractions of RTEL1 HHD2 were concentrated up to ∼7 mg/ml and used in the crystallization trials. Initial screening was conducted using Crystal Screen HT (Hampton Research; Cat. No. HR2-130) in 72-well oil immersion plates. The rod-like crystals of HHD2 appeared within two days at 4°C in many conditions of Crystal Screen as well as in the native crystallization buffer (20 mM Tris-HCl pH 7.4 at 25°C, 250 mM NaCl, 5% Glycerol, 2 mM DTT, 0.02% NaN_3_) of HHD2. Later, the HHD2 crystals were grown in the native buffer condition using the hanging drop vapour diffusion method.

The X-ray diffraction data sets were collected at ESRF beamline ID23-1 at Grenoble (France) using the PILATUS 6M pixel-array detector (DECTRIS Ltd., Switzerland). High-resolution data set was collected at an energy of 12.7 keV (**λ**= 0.976 Å). The crystals diffracted up to a maximum resolution of 1.49 Å. The diffraction data sets were processed using XDSAPP (Sparta et al., 2016). The RTEL1 HHD2 structure was solved at 1.6 Å resolution by using the Molecular Replacement with Phaser (McCoy et al., 2007), employing coordinates of the Alpha fold model of RTEL1 HHD2 (AF-Q9NZ71-F1) (Jumper et al., 2021) as the search model. Coot (Emsley and Cowtan, 2004), PHENIX (Adams et al., 2010), and REFMAC5 (Murshudov et al., 2011) were used for iterative model building and refinement. The R_work_ and R_free_ of the final model are 0.182 and 0.211, respectively. The quality of the final model was assessed using Coot (Emsley and Cowtan, 2004), the Molprobity server (Chen et al., 2010), and the wwPDB validation server (Young et al., 2017). Structural figures were generated using UCSF Chimera (Pettersen et al., 2004). The data statistics are presented in Table 1.

### DNA sample preparation

The DNA oligos (ssDNA-22 and csDNA-22) were purchased from Sigma-Aldrich in the lyophilized form. Both oligos are 22-mer ssDNA having complementary sequences to each other (Supplementary Table S2). The stock solutions of DNA samples were prepared by dissolving the lyophilized oligos separately in the required amount of NMR buffer (20 mM Tris-HCl pH 7.4 at 25°C, 50 mM NaCl, 2 mM DTT, 0.02% NaN_3_), and ITC buffer (10 mM Potassium phosphate pH 6.5 at 25°C, 50 mM KCl, 0.02% NaN_3_) for performing the NMR and ITC titrations, respectively. ssDNA-22 sample was used in the case of titration experiments with ssDNA. Samples of dsDNA-22 were made by mixing the equimolar amount of both the oligos, followed by incubation at 95°C for 10 min. The sample was finally reannealed by snap cooling on ice for 1 h.

### Solution NMR spectroscopy

All NMR spectra (2D ^1^H-^15^N HSQC and 2D ^1^H-^15^N TROSY HSQC) were acquired at 25°C on a Bruker 700 MHz spectrometer equipped with a cryogenic probe or a room temperature probe. Uniformly ^15^N-labeled protein samples were prepared in the NMR buffer (20 mM Tris-HCl pH 7.4 at 25°C, 50 mM NaCl, 2 mM DTT, 0.02% NaN_3_). 10% D_2_O (v/v) (Cambridge Isotope Laboratories; Cat. No. DLM-4-25) was added to the sample for the spectrometer deuterium lock. Typically, for the titration experiments, protein sample concentration was kept at 150 μM (in case of cryoprobe) or 250 μM (room temperature probe).

The NMR data were processed using Bruker Topspin and analyzed using the NMRFAM-SPARKY software (Lee et al., 2015).

The chemical shift perturbations (CSPs) were analyzed as combined amide chemical shift changes with equation 1: Δδ_NH_ (ppm) = [(Δδ^1^H)^2^ + (Δδ^15^N/5)^2^]^1/2^, where the chemical shift changes in the ^1^H and ^15^N dimensions are denoted by Δδ^1^H and Δδ^15^N respectively.

### Isothermal titration calorimetry (ITC)

Isothermal titration calorimetry (ITC) experiments were performed using a VP-ITC instrument (MicroCal, USA) at 25°C. The protein and DNA samples were prepared in ITC buffer (10 mM Potassium phosphate pH 6.5 at 25°C, 50 mM KCl, 0.02% NaN_3_) and thoroughly degassed before experiments.

The sample cell was filled with 10 μM of the DNA and titrated with 300 μM of the RTEL1 HHD2. 30 injections of the titrant (10 μl per injection) were performed with a spacing of 3 min between each injection. A control experiment (titration of HHD2 into the buffer; Figure 7C) was performed to know the heat of dilution of protein into the buffer. The integrated heat data was fitted for the one-site binding model using Origin software provided by the manufacturer. The Thermodynamic parameters of HHD2 and DNA interactions are presented in Table 2.

### Bioinformatic analysis

The disordered linker region between HHD1 and HHD2 domain was predicted through VSL2B (Obradovic et al., 2005) and IUPred3 (Erdős et al., 2021) servers. Multalign (Corpet, 1988) tool was used to perform the sequence alignment of the RTEL1 HHD2 domain from different vertebrate species. 32C binding peptide sequences were aligned with the help of Clustal Omega (Sievers et al., 2011). UCSF Chimera (Pettersen et al., 2004) was used for performing the secondary structure-based sequence alignment of different HHDs.

### HADDOCK Modelling

The NMR chemical shift perturbations (CSPs) data-driven docking of RTEL1 HHD2 and RPA 32C domain was performed through HADDOCK2.2 webserver with easy interface (van Zundert et al., 2016). In the case of HHD2 domain A1059, V1060, S1061, A1062, Y1063, L1064, A1065, D1066, A1067, R1068, R1069, G1075, S1077, Q1078, L1079, L1080, A1081, A1082, T1084, K1087, D1090, and D1134 were selected as active residues, and for 32C domain E223, G224, F227, I242, E252, G253, H254, Y256, T258, V259, D260, D261, D262, H263, F264, S266, T267, D268, and A269 were selected as active residues based on their perturbations above-average CSPs. Apart from these residues, other surrounding residues were considered passive residues.

Here, HADDOCK clustered 153 structures in 9 cluster(s), which represents 76 % of the water-refined models HADDOCK generated (Supplementary Table S1). The maximum number of models considered for clustering is 200.

After docking, the best-docked clusters are ranked according to their HADDOCK score. The ranking of the clusters is based on the average score of the top 4 members of each cluster. The score of the water-refined model is calculated as:

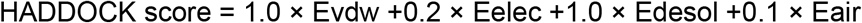

Where Evdw = intermolecular Van der Waals energy, Eelec = intermolecular electrostatic energy, Edesol = empirical desolvation energy, Eair = Restrain energy

The Z-score of a cluster indicates how many standard deviations from the average this cluster is in terms of score. The more negative HADDOCK score and Z-score of a cluster indicate better quality.

### Quantification and Statistical analysis

Quantification of co-immunoprecipitation data was performed through ImageJ software (Schneider et al., 2012). The quantified value and number of independent experiments are mentioned in the corresponding figure legend.

## ABBREVIATIONS

RTEL1: regulator of telomere elongation helicase 1
HHD: harmonin homology domain
RPA: replication protein A
NMR: nuclear magnetic resonance
ITC: isothermal titration calorimetry
CSP: chemical shift perturbation

## DATA AVAILABILITY

The atomic coordinates and structure factors for the RTEL1 HHD2 have been deposited in the Protein Data Bank under PDB accession code 7WU8.

## SUPPLEMENTARY DATA

Supplementary data are provided in the SI file.

## ACKNOWLEDGEMENTS

The authors acknowledge the Department of Science and Technology (DST) and Department of Biotechnology (DBT), India, for the NMR, ITC, and Confocal Microscopy facilities at the Indian Institute of Science, Bengaluru. The authors acknowledge funding for infrastructural support from the following programs of the Government of India: DST-FIST, UGC-CAS, and the DBT-IISc partnership program. The authors greatly acknowledge the provision of beamtime at the European Synchrotron Radiation Facility (ESRF), France. Authors acknowledge the X-ray diffraction facility for macromolecular crystallography at the Indian Institute of Science, Bengaluru, used for screening purposes, which is supported by the Science and Engineering Research Board, DST (DST-SERB) grant IR/SO/LU/0003/2010-PHASE-II. N.K. is a recipient of the GATE fellowship from the Ministry of Education, India. The authors thank Shuvra Shekhar Roy for his help in the initial purification of HHD1+2. The authors thank Dr. Mohsen Sarikhani for helping in the initial co-immunoprecipitation experiments. The authors thank Dr. Nukathoti Sivaji for his help in initial crystal screening and collecting the diffraction dataset at a home X-ray source.

## AUTHOR CONTRIBUTIONS

M.S. conceived the research; M.S. and N.K. designed the experiments. N.K. performed molecular cloning, protein purification, NMR, and ITC titration experiments. M.G. performed cloning, initial purification, and initial acquisition of the NMR spectrum of HHD1+2. N.K. and U.R. performed crystallization experiments; N.K., U.R., and M.S. solved and analyzed the X-ray structure. A.T., N.K., and N. R. S. performed the co-immunoprecipitation and immunofluorescence microscopy experiments. M.S. and N.K. wrote the manuscript with inputs from all other authors; all authors reviewed the final manuscript.

---

**Figure S1.**
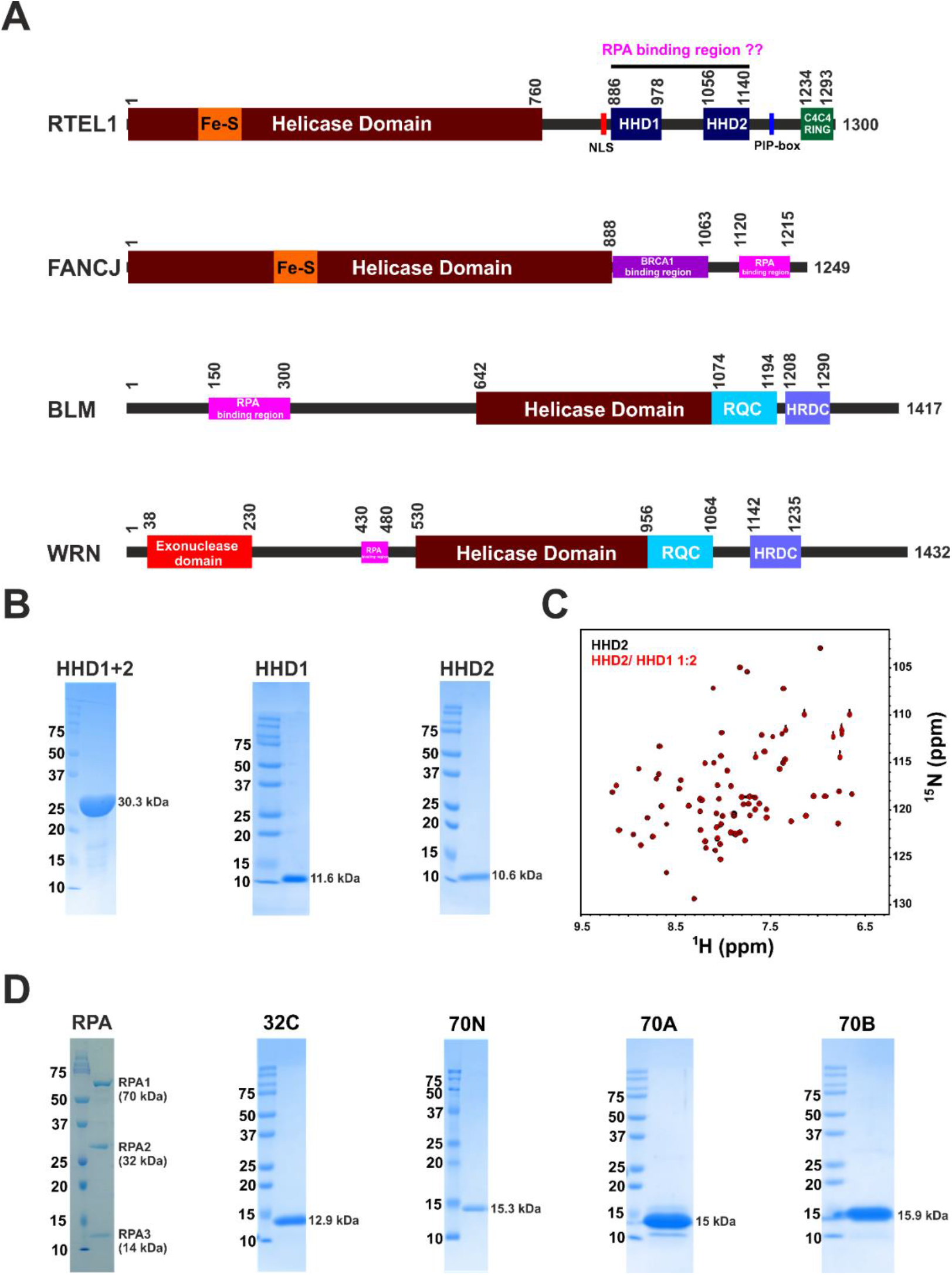
**(A)** Schematic of the domain organization of human RTEL1, FANCJ, BLM, and WRN helicases. The putative RPA binding region in RTEL1 is marked along with the known RPA binding region (pink rectangular box) in FANCJ, BLM, and WRN helicases. **(B)** SDS-PAGE gels of purified HHD1+2, HHD1, and HHD2. **(C)** Overlay of ^1^H -^15^N HSQC spectra of ^15^N-labelled HHD2 in the absence (black) and presence (red) of HHD1 (at 1:2 molar ratio). **(D)** SDS-PAGE gels of purified heterotrimeric complex of RPA and its different domains 32C, 70N, 70A, and 70B.

**Figure S2.**
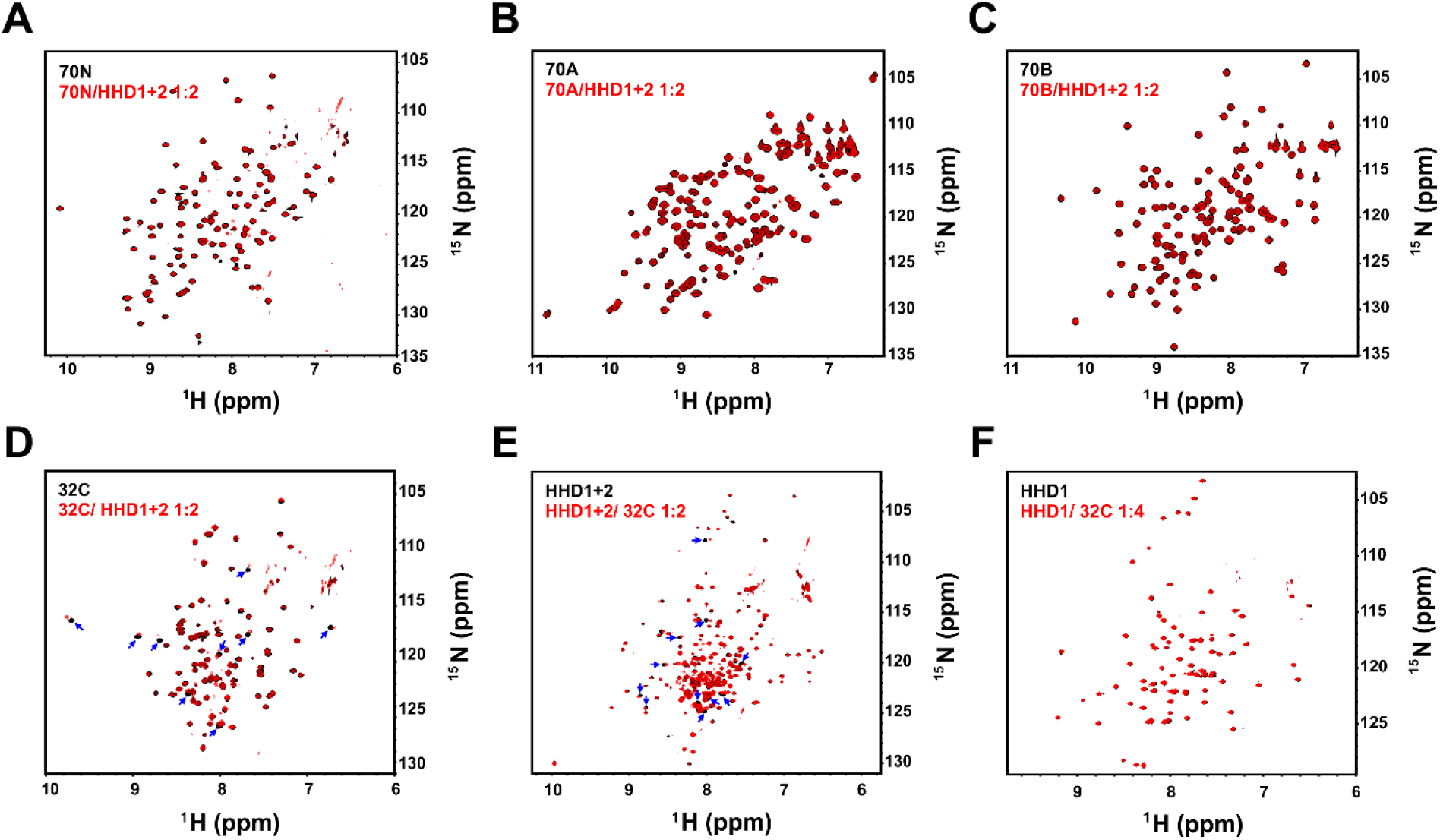
**(A)** Overlay of ^1^H -^15^N TROSY HSQC spectra of ^15^N-labelled 70N in the absence (black) and presence (red) of HHD1+2 (at 1:2 molar ratio). **(B)** Overlay of ^1^H -^15^N HSQC spectra of ^15^N-labelled 70A in the absence (black) and presence (red) of HHD1+2 (at 1:2 molar ratio). **(C)** Overlay of ^1^H -^15^N HSQC spectra of ^15^N-labelled 70B in the absence (black) and presence (red) of HHD1+2 (at 1:2 molar ratio). **(D)** Overlay of ^1^H -^15^N TROSY HSQC spectra of ^15^N-labelled 32C in the absence (black) and presence (red) of HHD1+2 (at 1:2 molar ratio). Residues with large CSPs are marked (blue arrows). **(E)** Overlay of ^1^H-^15^N TROSY HSQC spectra of ^15^N-labelled HHD1+2 in the absence (black) and presence (red) of 32C (at 1:2 molar ratio). Residues with large CSPs are marked (blue arrows). **(F)** Overlay of ^1^H-^15^N TROSY HSQC spectra of ^15^N-labelled HHD1 in the absence (black) and presence (red) of 32C (at 1:4 molar ratio). No significant CSPs were observed.

**Figure S3.**
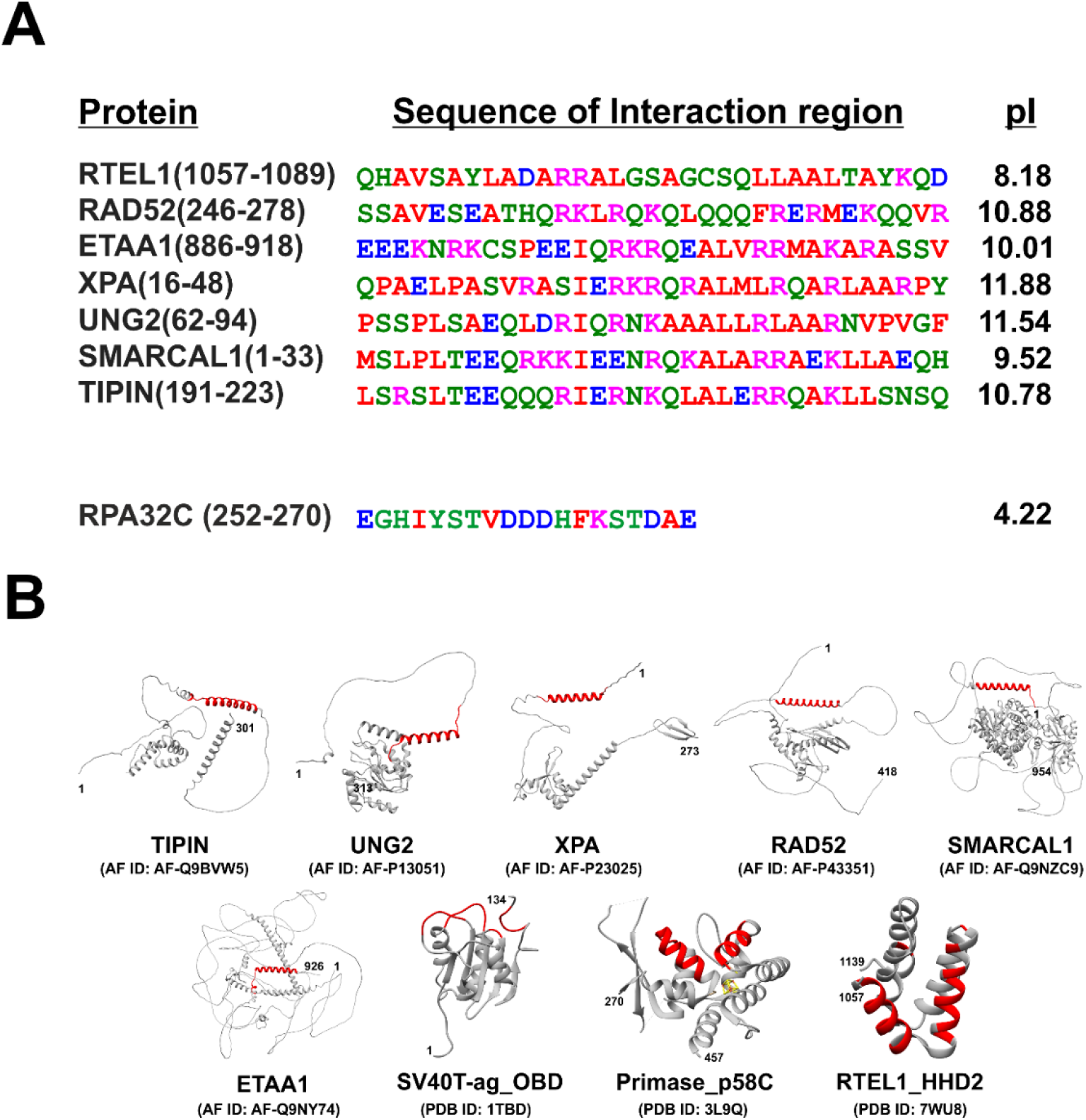
**(A)** Multiple sequence alignment of RPA 32C-interacting region in different proteins involved in DNA repair and replication. Sequence alignment was performed through Clustal Omega and further curated manually for analysis. As per the alignment, there is sequence similarity among ETAA1, XPA, UNG2, SMARCAL1, and TIPIN. Sequence boundaries and theoretical pI values of the sequences are indicated. The sequence corresponding to the complementary interacting region of RPA 32C is depicted at the bottom. Residue color code is based on their physicochemical properties (Clustal Omega). **(B)** RPA 32C-interacting helical regions are marked (in red) on the Alpha fold model structure of TIPIN, UNG2, XPA, RAD52, SMARCAL1, and ETAA1. In the case of origin binding domain (OBD) of SV40 T-antigen, C-terminal domain of primase p58, and HHD2 domain of human RTEL1, the RPA 32C binding surface is a more complex, comprising multiple helices and loops. PDB IDs and Alpha fold model IDs are indicated below each structure.

**Figure S4.**
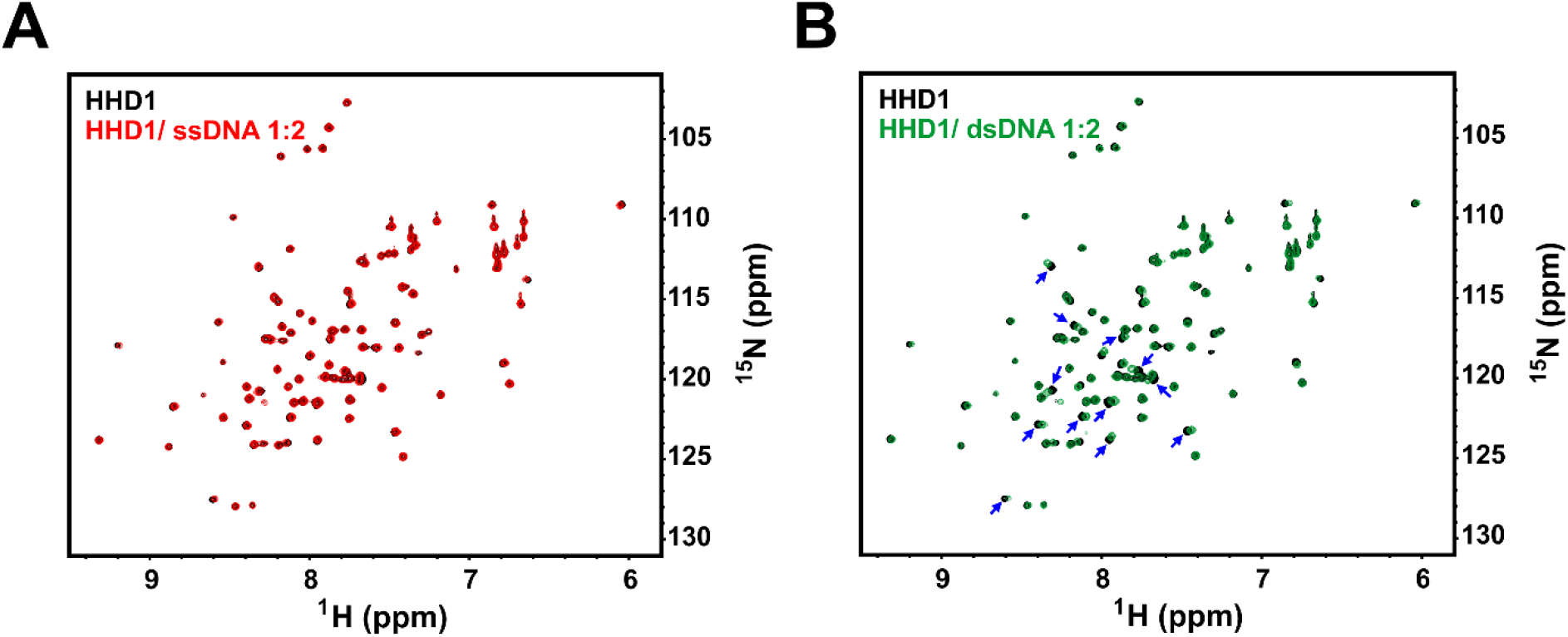
**(A)** Overlay of ^1^H-^15^N HSQC spectra of ^15^N-labeled HHD1 in the absence (black) and presence (red) of ssDNA-22. No significant CSPs were observed. **(B)** Overlay of ^1^H -^15^N HSQC spectra of ^15^N-HHD1 in the absence (black) and presence (green) of dsDNA-22 (at 1:2 molar ratio). Residues that showed CSPs are marked (blue arrow).

**Figure S5.**
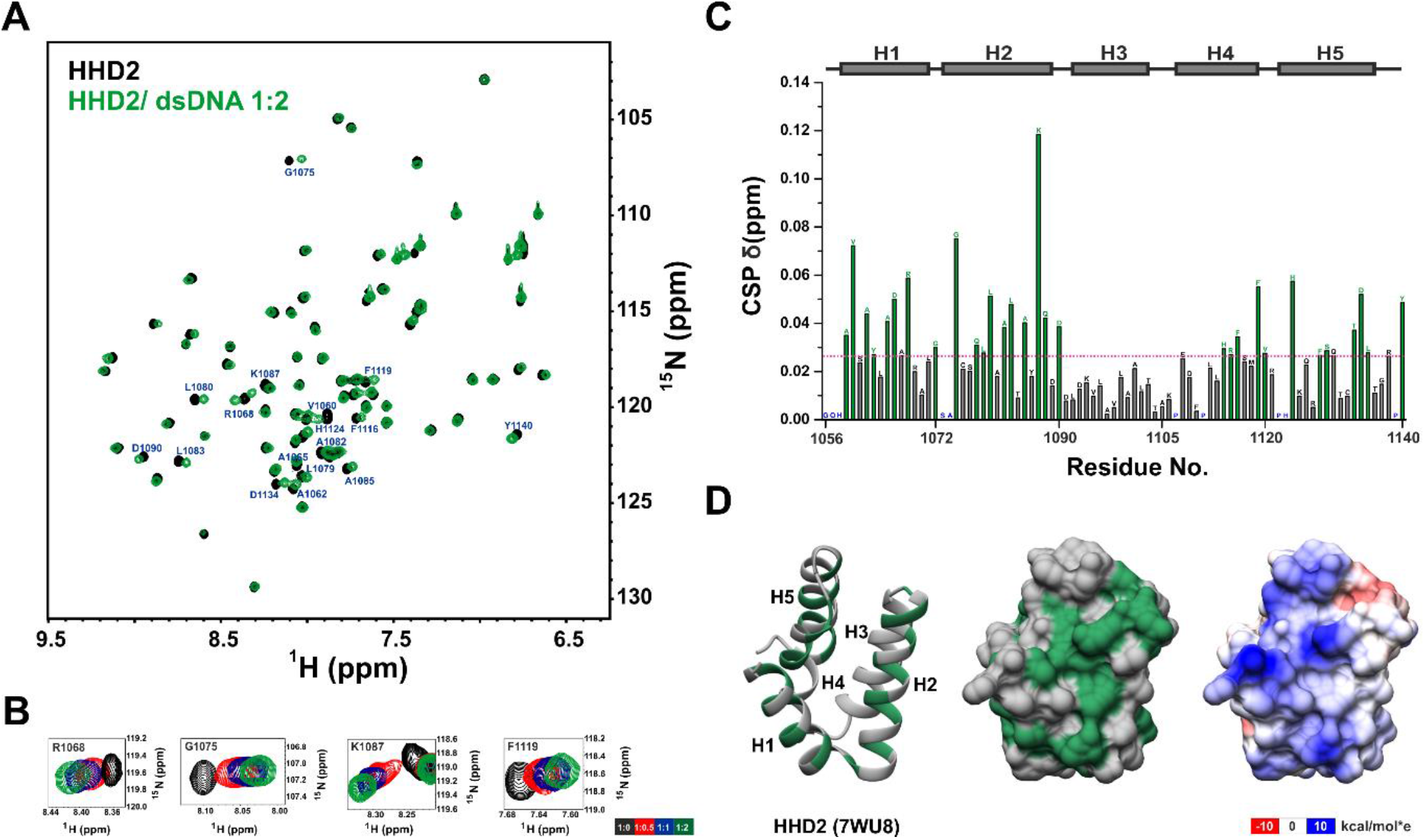
**(A)** Overlay of ^1^H-^15^N HSQC spectra of ^15^N-labeled HHD2 in the absence (black) and presence (green) of dsDNA-22 (at 1:2 molar ratio). Residues with large CSPs are labeled (in blue). **(B)** ^1^H-^15^N cross-peaks trajectory of representative residues R1068, G1075, K1087, and F1119 of HHD2 upon titration with dsDNA-22 at indicated molar ratios. **(C)** Quantification of CSPs in HHD2 upon titration with dsDNA-22 (at 1:2 molar ratio). Residues with more than average CSP (pink dash line) are marked as green bars and considered as significantly perturbed residues. Prolines and unassigned residues are colored in blue. The secondary structure corresponding to the HHD2 sequence is shown at the top. **(D)** Significantly perturbed residues (green) are marked on the structure of HHD2 (ribbon – left and surface – middle form). Most of these residues lie on the positively charged surface (right) of HHD2.

**Figure S6.**
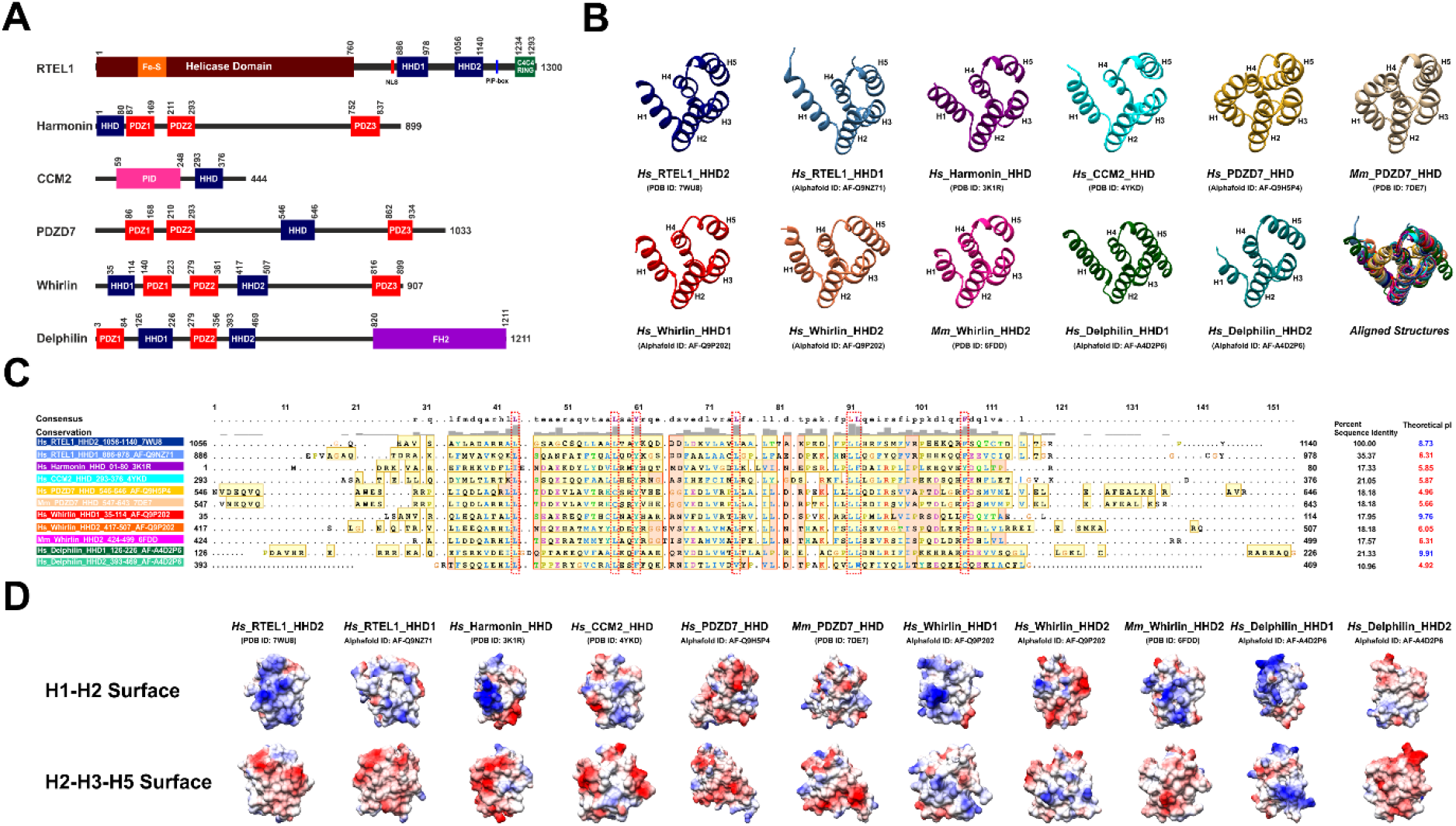
**(A)** Schematic of the domain organization of harmonin homology domains (HHDs) containing human proteins-RTEL1, Harmonin, CCM2, PDZD7, Whirlin, and Delphilin. Protein length and domain boundaries are mentioned. The HHDs are depicted as blue rectangles. **(B)** Experimental and model structures of all HHDs from humans (*Homo sapiens, Hs*) and two HHDs of the mouse (*Mus musculus, Mm*) are shown. PDB ID or Alpha fold model ID is indicated below each structure. The aligned structure of the HHDs is shown (bottom right). **(C)** Secondary structure-based multiple sequence alignment of RTEL1 HHD2 with HHDs present in other human and mouse proteins. The sequences forming secondary structures are highlighted in yellow. The consensus sequence and residue conservation bar are shown at the top. The extent of residue conservation is represented by the height of the conservation bar. Structurally conserved residues are marked with a red box. Percent sequence identity (with respect to RTEL1 HHD2) and theoretical pIs are shown on the right. RTEL1 HHD2, Whirlin HHD1, and Delphilin HHD1 have basic pI values (pI > 7, depicted in blue), whereas other HHDs have acidic pI values (pI < 7, depicted in red). Analyzing theoretical pIs indicates that the proteins with two HHDs (RTEL1, Whirlin, and Delphilin) have one HHD with acidic pI while another HHD with basic pI. **(D)** Comparison of the surface electrostatic potential of different HHDs. Two surfaces are shown. Only the HHD2 domain of RTEL1 shows a distinct positively charged surface (formed by helix H1 and H2) and negatively charged surface (formed by helix H2, H3, and H4).

**Table S1.** HADDOCK parameters of RTEL1 HHD2 – RPA 32C complex.

| S. No. | Parameters | Cluster 1 | Cluster 2 | Cluster 3 | Cluster 4 | Cluster 5 | Cluster 6 | Cluster 7 | Cluster 8 | Cluster 9 |
| --- | --- | --- | --- | --- | --- | --- | --- | --- | --- | --- |
| 1. | HADDOCK score | $-78.6 \pm 3.0$ | $-54.5 \pm 3.1$ | $-60.3 \pm 9.2$ | $-57.9 \pm 2.5$ | $-67.3 \pm 3.5$ | $-32.7 \pm 7.4$ | $-42.1 \pm 8.0$ | $-47.7 \pm 11.6$ | $-51.1 \pm 2.8$ |
| 2. | Cluster size | 79 | 22 | 13 | 12 | 11 | 4 | 4 | 4 | 4 |
| 3. | RMSD from the overall lowest-energy structure ($\text{\AA}$) | $0.7 \pm 0.4$ | $11.1 \pm 1.3$ | $2.4 \pm 1.1$ | $12.5 \pm 0.1$ | $5.1 \pm 0.4$ | $9.7 \pm 0.6$ | $3.7 \pm 0.2$ | $10.1 \pm 0.6$ | $4.0 \pm 0.7$ |
| 4. | Van der Waals energy ($\text{kcal mol}^{-1}$) | $-32.2 \pm 3.5$ | $-24.6 \pm 4.5$ | $-32.6 \pm 6.8$ | $-30.9 \pm 1.5$ | $-41.3 \pm 4.0$ | $-21.9 \pm 4.8$ | $-22.0 \pm 5.1$ | $-32.3 \pm 4.2$ | $-31.0 \pm 3.1$ |
| 5. | Electrostatic energy ($\text{kcal mol}^{-1}$) | $-317.4 \pm 25.0$ | $-221.2 \pm 19.3$ | $-205.8 \pm 18.4$ | $-188.8 \pm 17.8$ | $-200.3 \pm 14.3$ | $-155.7 \pm 31.7$ | $-235.4 \pm 19.8$ | $-137.4 \pm 51.8$ | $-163.5 \pm 22.9$ |
| 6. | Desolvation energy ($\text{kcal mol}^{-1}$) | $6.9 \pm 2.8$ | $2.2 \pm 1.9$ | $2.6 \pm 2.7$ | $-0.5 \pm 2.9$ | $3.5 \pm 2.3$ | $2.9 \pm 2.0$ | $9.6 \pm 3.1$ | $1.5 \pm 2.6$ | $1.2 \pm 3.9$ |
| 7. | Restraints violation energy ($\text{kcal mol}^{-1}$) | $102.6 \pm 25.3$ | $120.9 \pm 65.8$ | $108.1 \pm 64.7$ | $112.6 \pm 36.2$ | $106.3 \pm 38.4$ | $173.8 \pm 25.1$ | $174.7 \pm 26.5$ | $105.3 \pm 7.7$ | $113.4 \pm 16.5$ |
| 8. | Buried Surface Area ($\text{\AA}^2$) | $1322.7 \pm 29.4$ | $1087.2 \pm 36.7$ | $1190.6 \pm 80.1$ | $1087.0 \pm 86.4$ | $1260.2 \pm 34.8$ | $910.5 \pm 69.6$ | $1026.2 \pm 24.5$ | $1029.9 \pm 147.2$ | $1009.6 \pm 66.8$ |
| 9. | Z-score | -1.9 | 0.0 | -0.4 | -0.3 | -1.0 | 1.7 | 1.0 | 0.5 | 0.3 |

