## Supplementary Information for "Harmonin homology domain-mediated interaction of RTEL1 helicase with RPA and DNA provides mechanistic insight into its role in DNA repair"

#### SUPPLEMENTARY FIGURES

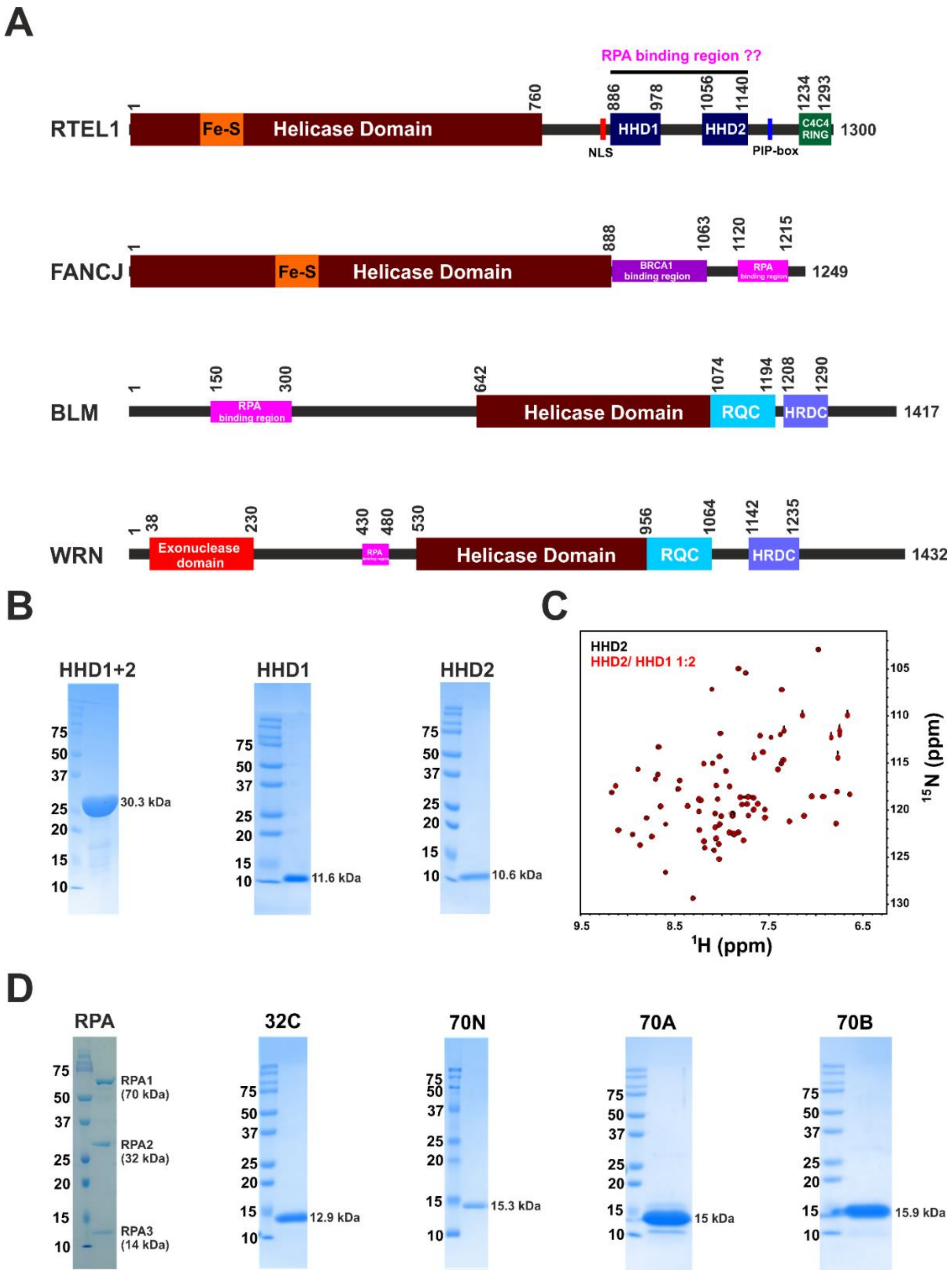


**Figure S1. (A)** Schematic of the domain organization of human RTEL1, FANCJ, BLM, and WRN helicases. The putative RPA binding region in RTEL1 is marked along with the known RPA binding region (pink rectangular box) in FANCJ, BLM, and WRN helicases (Kang et al., 2018; Yeom et al., 2019)**. (B)** SDS-PAGE gels of purified HHD1+2, HHD1, and HHD2**. (C)** Overlay of $^{1}$H -$^{15}$N HSQC spectra of $^{15}$N-labelled HHD2 in the absence (black) and presence (red) of HHD1 (at 1:2 molar ratio). **(D)** SDS-PAGE gels of purified heterotrimeric complex of RPA and its different domains 32C, 70N, 70A, and 70B.

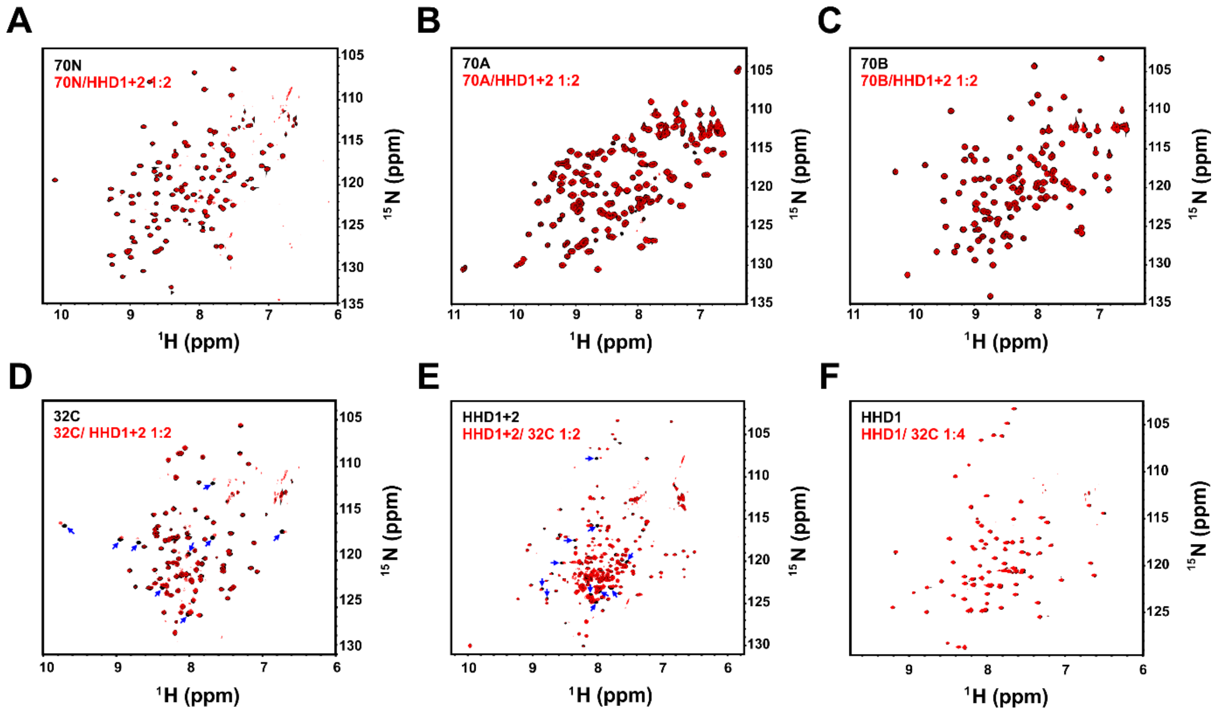


**Figure S2. (A)** Overlay of $^{1}$H -$^{15}$N TROSY HSQC spectra of $^{15}$N-labelled 70N in the absence (black) and presence (red) of HHD1+2 (at 1:2 molar ratio). **(B)** Overlay of $^{1}$H -$^{15}$N HSQC spectra of $^{15}$N-labelled 70A in the absence (black) and presence (red) of HHD1+2 (at 1:2 molar ratio). **(C)** Overlay of $^{1}$H -$^{15}$N HSQC spectra of $^{15}$N-labelled 70B in the absence (black) and presence (red) of HHD1+2 (at 1:2 molar ratio). **(D)** Overlay of $^{1}$H -$^{15}$N TROSY HSQC spectra of $^{15}$N-labelled 32C in the absence (black) and presence (red) of HHD1+2 (at 1:2 molar ratio). Residues with large CSPs are marked (blue arrows). **(E)** Overlay of $^{1}$H-$^{15}$N TROSY HSQC spectra of $^{15}$N-labelled HHD1+2 in the absence (black) and presence (red) of 32C (at 1:2 molar ratio). Residues with large CSPs are marked (blue arrows). **(F)** Overlay of $^{1}$H-$^{15}$N TROSY HSQC spectra of $^{15}$N-labelled HHD1 in the absence (black) and presence (red) of 32C (at 1:4 molar ratio). No significant CSPs were observed.

**A**

| Protein | Sequence of Interaction region | pI |
|---|---|---|
| RTEL1(1057-1089) | QHAVSAYLADARRALGSAGCSQLLAALTAYKQD | 8.18 |
| RAD52(246-278) | SSAVESEATHQRKLRQKQLQQQFRERMEKQQVR | 10.88 |
| ETAA1(886-918) | EEEKNRKCSPEEIQRKRQEALVRRMAKARASSV | 10.01 |
| XPA(16-48) | QPAELPASVRASIERKRQRALMLRQARLAARPY | 11.88 |
| UNG2(62-94) | PSSPLSAEQLDRIQRNKAAALLRLAARNVPVGF | 11.54 |
| SMARCAL1(1-33) | MSLPLTEEQRKKIEENRQKALARRAEKLLAEQH | 9.52 |
| TIPIN(191-223) | LSRSLTEEQQQRIERNKQLALERRQAKLLSNSQ | 10.78 |
| RPA32C (252-270) | EGHIYSTVDDDHFKSTDAE | 4.22 |

**B**

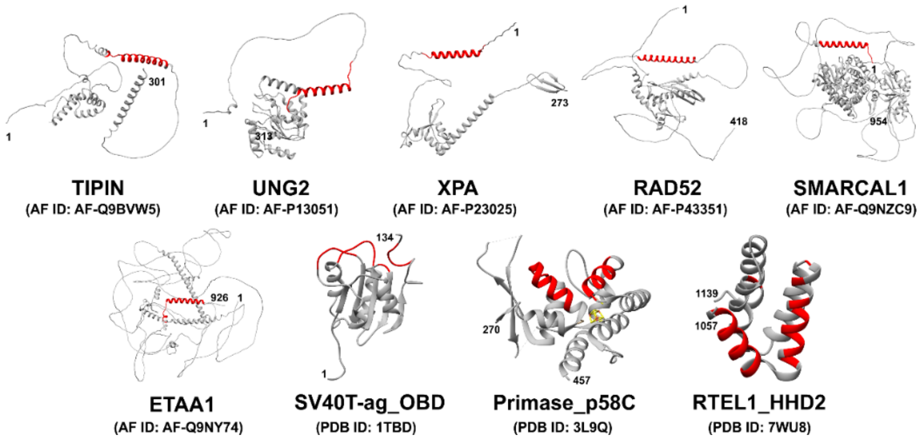


**Figure S3. (A)** Multiple sequence alignment of RPA 32C-interacting region in different proteins involved in DNA repair and replication (Ali et al., 2010; Bass et al., 2016; Feldkamp et al., 2014; Mer et al., 2000; Xie et al., 2014). Sequence alignment was performed through Clustal Omega and further curated manually for analysis. As per the alignment, there is sequence similarity among ETAA1, XPA, UNG2, SMARCAL1, and TIPIN. Sequence boundaries and theoretical pI values of the sequences are indicated. The sequence corresponding to the complementary interacting region of RPA 32C is depicted at the bottom. Residue color code is based on their physicochemical properties (Clustal Omega). **(B)** RPA 32C-interacting helical regions are marked (in red) on the Alpha fold model structure of TIPIN, UNG2, XPA, RAD52, SMARCAL1, and ETAA1. In the case of origin binding domain (OBD) of SV40 T-antigen, C-terminal domain of primase p58, and HHD2 domain of human RTEL1, the RPA 32C binding surface is a more complex, comprising multiple helices and loops (Arunkumar et al., 2005;

Vaithiyalingam et al., 2010). PDB IDs and Alpha fold model IDs are indicated below each structure.

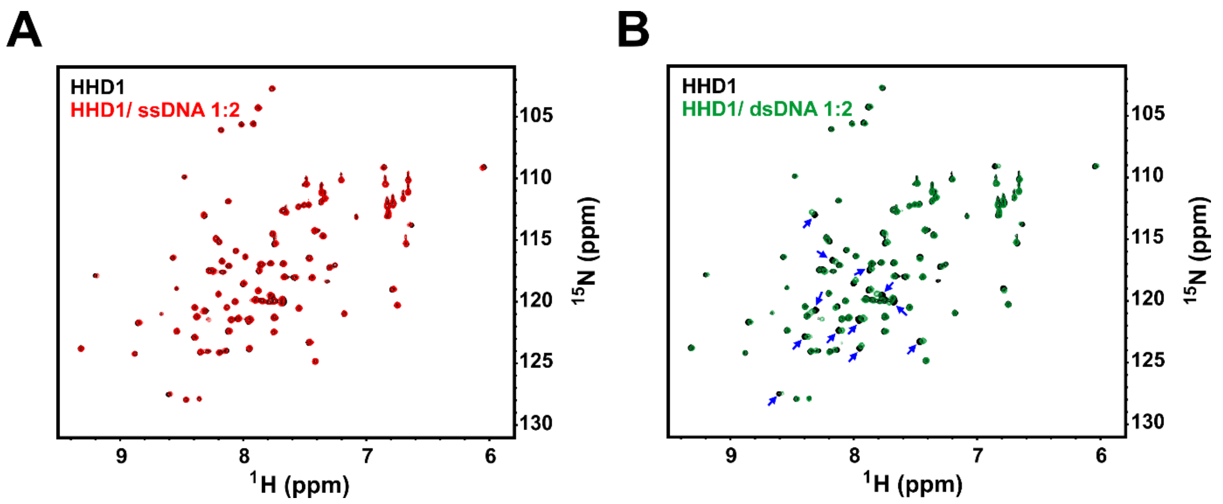


**Figure S4. (A)** Overlay of $^{1}$H-$^{15}$N HSQC spectra of $^{15}$N-labeled HHD1 in the absence (black) and presence (red) of ssDNA-22. No significant CSPs were observed. **(B)** Overlay of $^{1}$H -$^{15}$N HSQC spectra of $^{15}$N-HHD1 in the absence (black) and presence (green) of dsDNA-22 (at 1:2 molar ratio). Residues that showed CSPs are marked (blue arrow).

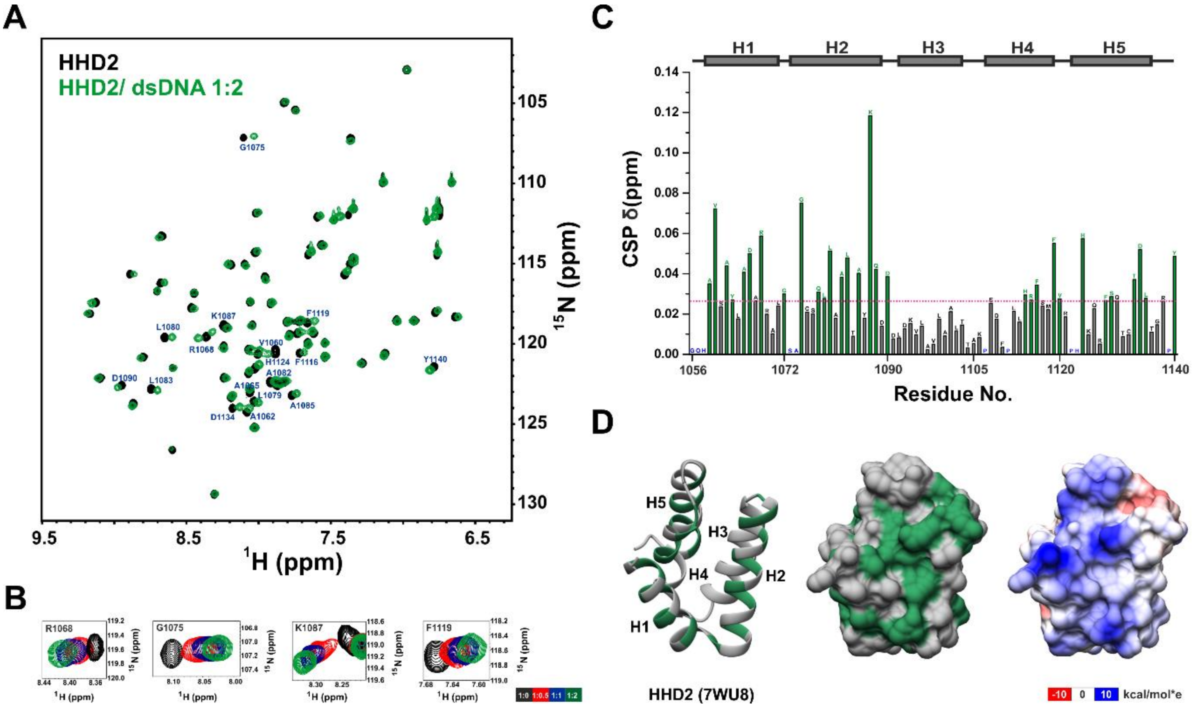


**Figure S5. (A)** Overlay of $^{1}$H-$^{15}$N HSQC spectra of $^{15}$N-labeled HHD2 in the absence (black) and presence (green) of dsDNA-22 (at 1:2 molar ratio). Residues with large CSPs are labeled (in blue). **(B)** $^{1}$H-$^{15}$N cross-peaks trajectory of representative residues R1068, G1075, K1087, and F1119 of HHD2 upon titration with dsDNA-22 at indicated molar ratios. **(C)** Quantification of CSPs in HHD2 upon titration with dsDNA-22 (at 1:2 molar ratio). Residues with more than average CSP (pink dash line) are marked as green bars and considered as significantly perturbed residues. Prolines and unassigned residues are colored in blue. The secondary structure corresponding to the HHD2 sequence is shown at the top. **(D)** Significantly perturbed residues (green) are marked on the structure of HHD2 (ribbon – left and surface – middle form). Most of these residues lie on the positively charged surface (right) of HHD2.

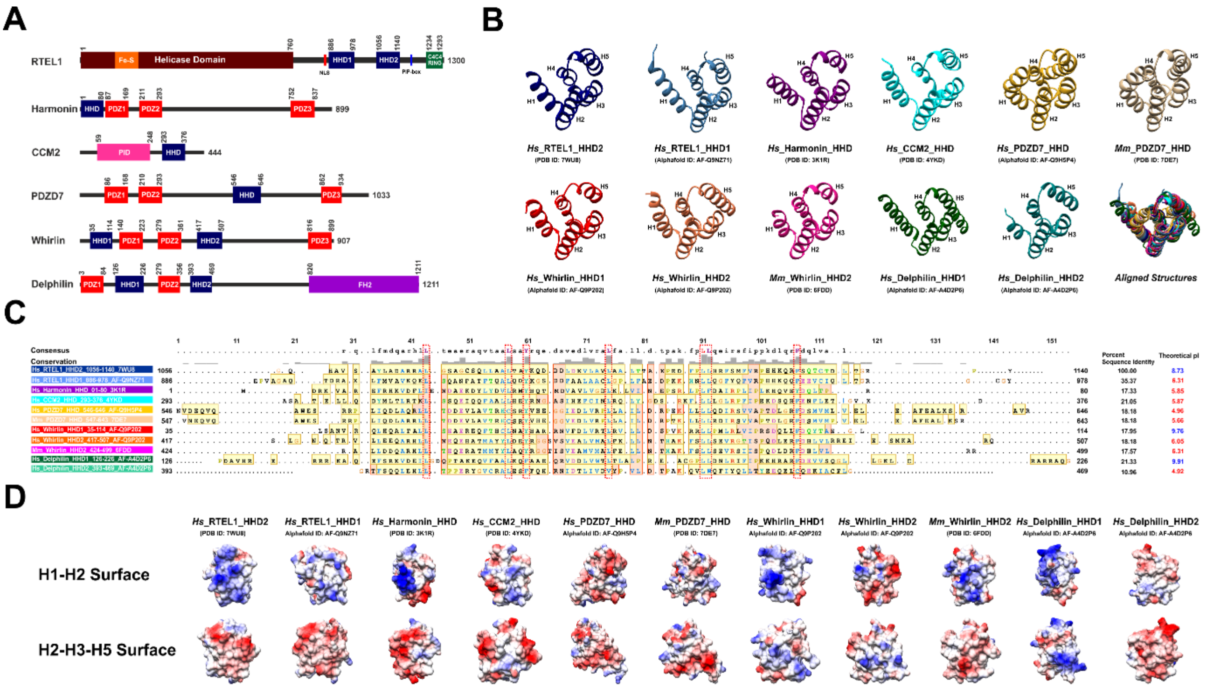


**Figure S6. (A)** Schematic of the domain organization of harmonin homology domains (HHDs) containing human proteins- RTEL1, Harmonin, CCM2, PDZD7, Whirlin, and Delphilin. Protein length and domain boundaries are mentioned. The HHDs are depicted as blue rectangles. **(B)** Experimental and model structures of all HHDs from humans (*Homo sapiens, Hs*) and two HHDs of the mouse (*Mus musculus, Mm*) are shown (Delhommel et al., 2018; Lin et al., 2021; Wang et al., 2015; Yan et al., 2010). PDB ID or Alpha fold model ID is indicated below each structure. The aligned structure of the HHDs is shown (bottom right). **(C)** Secondary structure-based multiple sequence alignment of RTEL1 HHD2 with HHDs present in other human and mouse proteins. The sequences forming secondary structures are highlighted in yellow. The consensus sequence and residue conservation bar are shown at the top. The extent of residue conservation is represented by the height of the conservation bar. Structurally conserved residues are marked with a red box. Percent sequence identity (with respect to RTEL1 HHD2) and theoretical pIs are shown on the right. RTEL1 HHD2, Whirlin HHD1, and Delphilin HHD1 have basic pI values (pI > 7, depicted in blue), whereas other HHDs have acidic pI values (pI < 7, depicted in red). Analyzing theoretical pIs indicates that the proteins with two HHDs (RTEL1, Whirlin, and Delphilin) have one HHD with acidic pI while another HHD with basic pI.

**(D)** Comparison of the surface electrostatic potential of different HHDs. Two surfaces are shown. Only the HHD2 domain of RTEL1 shows a distinct positively charged surface (formed by helix H1 and H2) and negatively charged surface (formed by helix H2, H3, and H4).

### SUPPLEMENTARY TABLES

#### Table S1. HADDOCK parameters of RTEL1 HHD2 – RPA 32C complex

| S. No. | Parameters | Cluster 1 | Cluster 2 | Cluster 3 | Cluster 4 | Cluster 5 | Cluster 6 | Cluster 7 | Cluster 8 | Cluster 9 |
|---|---|---|---|---|---|---|---|---|---|---|
| 1. | HADDOCK score | -78.6 ± 3.0 | -54.5 ± 3.1 | -60.3 ± 9.2 | -57.9 ± 2.5 | -67.3 ± 3.5 | -32.7 ± 7.4 | -42.1 ± 8.0 | -47.7 ± 11.6 | -51.1 ± 2.8 |
| 2. | Cluster size | 79 | 22 | 13 | 12 | 11 | 4 | 4 | 4 | 4 |
| 3. | RMSD from the overall lowest-energy structure (Å) | 0.7 ± 0.4 | 11.1 ± 1.3 | 2.4 ± 1.1 | 12.5 ± 0.1 | 5.1 ± 0.4 | 9.7 ± 0.6 | 3.7 ± 0.2 | 10.1 ± 0.6 | 4.0 ± 0.7 |
| 4. | Van der Waals energy (kcal $mol^{-1}$) | -32.2 ± 3.5 | -24.6 ± 4.5 | -32.6 ± 6.8 | -30.9 ± 1.5 | -41.3 ± 4.0 | -21.9 ± 4.8 | -22.0 ± 5.1 | -32.3 ± 4.2 | -31.0 ± 3.1 |
| 5. | Electrostatic energy (kcal $mol^{-1}$) | -317.4 ± 25.0 | -221.2 ± 19.3 | -205.8 ± 18.4 | -188.8 ±17.8 | -200.3 ± 14.3 | -155.7 ± 31.7 | -235.4 ± 19.8 | -137.4 ± 51.8 | -163.5 ± 22.9 |
| 6. | Desolvation energy (kcal $mol^{-1}$) | 6.9 ± 2.8 | 2.2 ± 1.9 | 2.6 ± 2.7 | -0.5 ± 2.9 | 3.5 ± 2.3 | 2.9 ± 2.0 | 9.6 ± 3.1 | 1.5 ± 2.6 | 1.2 ± 3.9 |
| 7. | Restraints violation energy (kcal $mol^{-1}$) | 102.6 ± 25.3 | 120.9 ± 65.8 | 108.1 ±64.7 | 112.6 ±36.2 | 106.3 ± 38.4 | 173.8 ± 25.1 | 174.7 ± 26.5 | 105.3 ± 7.7 | 113.4 ± 16.5 |
| 8. | Buried Surface Area ($Å^2$) | 1322.7 ± 29.4 | 1087.2 ± 36.7 | 1190.6 ± 80.1 | 1087.0 ± 86.4 | 1260.2 ± 34.8 | 910.5 ± 69.6 | 1026.2 ± 24.5 | 1029.9 ± 147.2 | 1009.6 ±66.8 |
| 9. | Z-score | -1.9 | 0.0 | -0.4 | -0.3 | -1.0 | 1.7 | 1.0 | 0.5 | 0.3 |

**Table S2. Oligonucleotides used in this study**

| S. No. | Oligonucleotides Name | Sequence (5’–3’) |
|---|---|---|
| | *Construction of plasmids for E. coli expression* | |
| 1. | HHD1+2_FP | GTCACATATGGAGCCCGTGG |
| 2. | HHD1+2_RP | GTACCTCGAGCTAGTAGGGCCGG |
| 3. | 70N_FP | GTCACATATGGTCGGCCAGCTGAGCGAG |
| 4. | 70N_RP | GTCAGGATCCTTATTCATTATAGGGCACTGG |
| 5. | 70A_FP | GTCACATATGCAGTCCAAAGTGGTGCCCATTG |
| 6. | 70A_RP | GTCAGGATCCTTAGTCCTCACAGGGCATGACGG |
| 7. | 70B_FP | GTCACATATGCAGTTTGATTTCACGGGGATTG |
| 8. | 70B_RP | GTCAGGATCCTTATAAGGCTTGTCCTTCTGCG |
| 9. | 32C_FP | GTCACATATGGCCAACAGCCAGCCCTCA |
| 10. | 32C_RP | GTCAGGATCCTTATTCTGCATCTGTGGATTT |
| | *DNA sample for ITC and NMR titrations* | |
| 11. | ssDNA-22 | AGGATTAGGATTAGGATTAGGA |
| 12. | csDNA-22 | TCCTAATCCTAATCCTAATCCT |

**Table S3. Plasmids used in this study**

| S. No. | Plasmids Name | Source | Identifier |
|---|---|---|---|
| 1. | pcDNA3.1+N-HA-RTEL1 | Genscript; This study | Clone ID OHu14298 |
| 2. | pcDNA3.1+N-HA-RTEL1-ΔHHD1+2 | Genscript; This study | N/A |
| 3. | p11d-tRPA (123) | Addgene (Henricksen et al., 1994) | Plasmid # 102613 |
| 4. | pET-28a (+) HHD1 | (Kumar et al., 2022) | N/A |
| **5.** | pET-28a (+) HHD2 | (Kumar et al., 2022) | N/A |
| **6.** | pET-28a (+) HHD1+2 | This study | N/A |
| **7.** | pET-15b-RPA 70N | This study | N/A |
| **8.** | pET-15b-RPA 70A | This study | N/A |
| **9.** | pET-15b-RPA 70B | This study | N/A |
| **10.** | pET-15b-RPA 32C | This study | N/A |
